# Telomeric G-quadruplex Intermediates unveiled by Complex Markov Network Analysis

**DOI:** 10.1101/2025.01.16.633340

**Authors:** A. Sáinz-Agost, F. Falo, A. Fiasconaro

## Abstract

G-quadruplexes are secondary, non-canonical RNA/DNA structures formed by guanine-rich sequences assembled into four-stranded helical structures by the progressive stacking of G-Tetrads, planar arrangements of guanines stabilised by monovalent ions such as K^+^ or Na^+^. Their stability plays a very important role in the prevention of DNA degradation, leading to the promotion or inhibition of specific biological pathways upon formation. In this work, we explore the occurrences of intermediates originating from the unfolding of these structures by using all-atom simulations, analyzing a small number of significant reaction coordinates to follow the evolution of the system by applying a mesoscopic simplification of the structures followed by two different dimensionality reduction techniques: Principal Component Analysis (PCA) and time-Independent Component Analysis (tICA). The data of the reduced trajectories are then encoded into a Complex Markov Network which, in conjunction with an Stochastic Steepest Descent, provides a hierarchical organization of the different nodes into basins of attraction. This procedure is able to reveal the main intermediates and the most relevant transitions the system undergoes in its denaturation path.

## 1 Introduction

G-quadruplexes (G4s) are secondary, non-canonical structures arising from either one or multiple guanine rich DNA or RNA chains, and are very common in the human genome^1,2^. These conformations emerge from the gradual stacking of *G-tetrads*, planar arrangements of four guanines stabilized by Hoogsteen hydrogen bonds^3–5^. The stacking process is further aided by the presence of either monovalent or divalent cations along the central channel these tetrads define^6^, with typically K^+^ or Na^+^ being involved.

The relevance and recent interest in these structures comes from their biological function. G-quadruplexes have an important role in DNA stability^7^ as well as in regulation processes^8–10^, the latter associated to either the over- or under-exposure of different binding sites of interest upon adoption of a G-quadruplex topology. The presence of these structures at the moment of DNA replication can lead to the interruption of this process, destabilizing the helicases^11^ and, combining with other factors, can lead to epigenetic instability^12^. Similarly, overabundance of guanine in mRNA can lead to difficulties in protein translation in the ribosome. Several works have investigated their gene promotion and repression capabilities^13,14^, as well as its relevance as a potential therapeutic binding site for cancer treatment, given that the telomeric regions of the chromosomes have a tendency to form these structures^15,16^.

G-quadruplexes can adopt diverse topologies^3^, primarily influenced by two key factors: the number of single-stranded DNA/RNA chains participating in their formation (from either a single or several molecules, the latter called intermolecular G4s) and the particular twists and turns present in their backbones, dividing them into parallel (all guanine strands oriented in the same direction), antiparallel (neighboring strands oriented in opposite directions) or hybrid. Our research focuses specifically on unimolecular quadruplexes, formed by a single stranded DNA (ssDNA) chain. Within this context, we examined one of the multiple structures available: the *parallel* G-quadruplex, which has all guanine tracts are oriented in the same direction, in which the loops connect the top of a guanine track with the bottom of the next. This conformation is depicted on Fig. 1.

**Figure 1.**
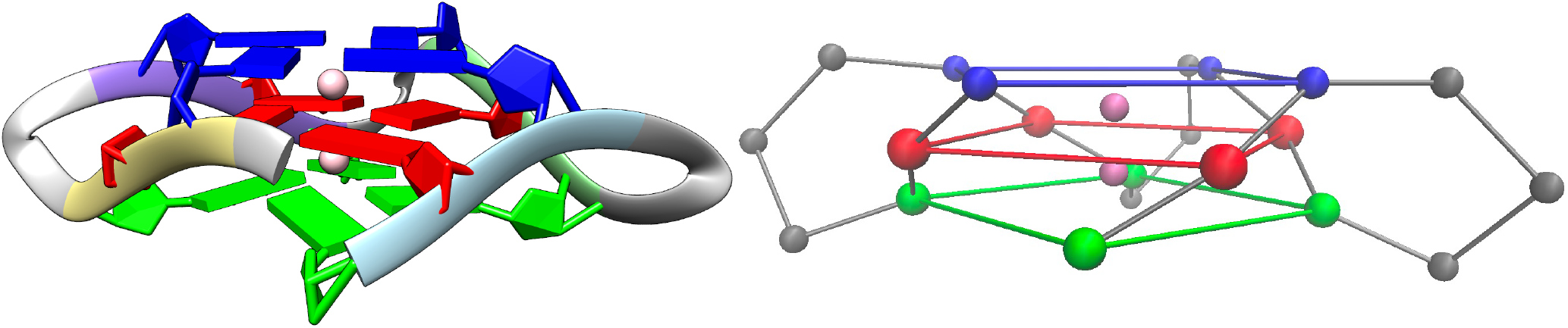
On the left, parallel conformation sourced from the PDB (1KF1, ribbon representation in Chimera). On the right, the resulting structure after the coarse graining procedure. The different G-tetrads on the structures are marked in different colors, the loops in grey and the ions in pink. As concerns the PDB structure, the different guanine tracts are also represented in homogeneous colors: yellow (first tract, 5^*′*^-end), purple (second tract), lime (third tract) and ice blue (fourth tract, 3^*′*^-end), respectively.

The mechanical stability of G-quadruplexes under force has been extensively documented^6,17–19^. Thus, the studies in this field have turned towards the study of their thermodynamical properties^20,21^. In particular, the observation and subsequent characterization of different conformations and folding intermediates of G-quadruplexes has become an object of large interest, due to the involvement of these structures in genetic regulation.^22^.

The goal of this work is to focus into the thermal unfolding of G-quadruplexes and investigate the presence of intermediate structures in the unfolding pathway. The stability of the G4 is affected by several factors both external, such as ionic concentration^23,24^, presence of other elements in the solvent^25,26^ and bath temperature, or internal, such as strand orientation (parallel, antiparallel or hybrid), length and structure of the loops^24,27,28^, and chain rigidity^18^.

Molecular Dynamics (MD) simulations have proven to be a valid and powerful tool in examining both the thermal and mechanical stability of biomolecular structures, also G-quadruplexes, revealing complex energy landscapes and characterizing their folding kinetics^29,30^. In the context of MD, we have used the *Replica Exchange Molecular Dynamics* (REMD)^31,32^ method to study the evolution of our system through the parallel simulation of 8 copies of the human parallel G-quadruplex (PDB: 1KF1)^33^ in a temperature range close to its reported melting point (65^*°*^), observing an unfolding event in only one of the eight replicas. To efficiently analyze the resulting high-dimensional data, a coarse-graining approach based on a consolidated mesoscopic model^17,18^ was applied, followed by the dimensionality reduction techniques, specifically Principal Component Analysis (PCA)^34,35^ and time-lagged Independent Component Analysis (tICA)^36^. These techniques allowed us to reduce the complexity of the data and retain only the most relevant information about the system. Using Complex Markov Networks (CMN)^37–40^ combined with a stochastic steepest descent algorithm, we constructed a series of networks describing the unfolding process. These networks outlined the main conformations the system adopts as it unfolds, identifying both stable and transitory intermediates. The description provided by tICA proved to be clearer than that of PCA, sometimes identifying a temporal evolution of the denaturation pathway.

This work is structured as follows: in Section 2 we elaborate on the simulation setup, the different dimensionality reduction approaches, and the description and construction of the Complex Markov Networks. Section 3 contains a discussion of the simulation findings, the effect of the presence of the cations inside the G4 structure on the unfolding dynamics, the results arising from the application of the two dimensionality reduction procedures considered, as well as the resulting CMNs describing the unfolding. The final conclusions are contained in Section 4. Additional information regarding both methods and results can be found in the Supplementary Material associated to this publication.

## 2 Methods

### 2.1 All-atom simulations with Gromacs

The initial structure for the simulation, the intramolecular parallel DNA G-quadruplex (1KF1)^33^, formed by GGGTTA repeat units, as 5^*′*^-AGGGTTAGGGTTAGGGTTAGGG-3^*′*^, was retrieved from the Protein Data Bank (PDB).

The simulations were carried out using the software Gromacs^41,42^, with the help of the force field amber modification Parmbsc0^43^, which has been used for both DNA^44^ and G-quadruplex analysis^6^ and has been proven to be the best option for our calculations^45^, although other options, such as OL15^46^ or Parmbsc1^47^, could also be considered. Control measures to verify the stability of the complexes in normal conditions under the force field of interest were carried out, obtaining the expected results.

The structures were solvated using the Tip3P water model^48^ in a periodic cubic box 1 nm larger than the diameter of the DNA body. Afterwards, more potassium K^+^ ions were introduced in the simulation box to counteract the negative charges present in the DNA chains, resulting in an average potassium concentration of 0.14 M, with no negative ions added to the mix.

The energy of the solvent was minimized using a steepest descent algorithm for up to 50000 iterations, leaving the G4s frozen. This was followed by equilibration in the NVT and NPT ensembles, both lasting for 100 ps using a leap-frog integrator, with a Berendsen thermostat and a Parrinello-Rahman barostat. The electrostatic interactions were treated via the particle mesh Ewald method, establishing its cutoff, as well as the one for the Lennard-Jones interactions, at 1 nm. All simulations performed used a time step of 2 fs, and simulation data was recorded every 20 ps.

To observe the unfolding intermediates of the structures, our simulations have been performed at temperatures near the melting point of the parallel G4, being slightly higher than 65°C^49^. At these temperatures, the unfolding kinetics are notoriously slow.

To increase the unfolding probability of the G4s and avoid that the system may get stuck in a metastable state^50^, we applied in our simulations the *Replica Exchange Molecular Dynamics* (REMD) method, also called Parallel Tempering^31,32^. This methodology consists in running concurrent simulations of multiple copies of the entire system (structure, solvent and ions), each of them at different temperatures. We refer to the simulation boxes containing a copy of the system at a given temperature as *replicas*. Once the simulation is started, after a sufficient time lapse (below described)^51–53^, attempts are made to exchange the content between replicas at different temperatures, re-scaling the velocities of the particles in the process. The probability of such exchanges is determined by a prescribed algorithm, in our case the Metropolis criterion^54^. This interchange of conformations at diverse temperatures aids the system in overcoming high energy barriers.

We constructed eight replicas, with assigned temperatures exceeding the nominal melting point of the structures (65°C). This adjustment was made in consideration of the ionic strength found in our simulations, which is higher than that of the experimental measurements, a fact that has been shown to increase the melting temperature T_m_^23^. The concrete values were in the interval [343, 345, …, 357] K.

As refers to the exchange time between replicas, we followed a different criterion with respect to the standard procedure^51–53^. In fact, in order to guarantee the subsequent PCA and tICA analysis under a well determined temperature, we have used a thermalization condition after each replica exchange. This way, the minimum switching time has been set to a time interval of 200ns, time during which the energy potential can be considered thermalized, according to the relaxation of the potential energy visible in the Supplementary Material. The consequence of this choice lays in a reduced unfolding probability, but guarantees the correctness of the PCA and tICA procedure.

### 2.2 Mesoscopic discretization of the model

The aim of this work is to extract the main conformations the human parallel G-quadruplex adopts along its unfolding trajectory. The method employed for the analysis relies partially in the diagonalization of different correlation matrices, whose dimension corresponds to the number of coordinates involved in the description of the all-atom trajectories of these structures, as explained in later paragraphs. In order to reduce the size of these matrices as well as to only focus on the conformations that lead to significant changes in the backbone of the G-quadruplexes, we decided to coarse-grain the structures before this analysis.

Specifically, the coarse-graining procedure, consistent with prior investigations in our research group^17,18^, involved the removal of water and ions from the simulation box, retaining only the DNA bases, which were subsequently reduced to unique identical beads. The positions of these beads were determined as the center of mass for the guanines and the position of the phosphates for the other nucleotides composing the loops. The transformation of gromacs coordinates into a coarse-grained system was executed using Python 3.9.18 in conjunction with the MDAnalysis package^55,56^. The resulting structure is depicted on the right side of Fig. 1.

### 2.3 Dimensionality reduction techniques

Following the coarse-graining, our system now comprises 21 beads, each characterized by its position along the three Cartesian axes, resulting in a total of 63 degrees of freedom. Extensive research^57,58^ has demonstrated that the majority of these degrees of freedom do not contribute significantly to our understanding of the system. Instead, the essential aspects can be effectively captured using a reduced set of reaction coordinates derived from combinations of the original ones.

In this study, we employ two distinct methods to diminish the system’s dimensionality while retaining a substantial amount of relevant information. These methodologies are recognized as “Principal Component Analysis” (PCA) and “time-Independent Component Analysis.” (tICA).

#### 2.3.1 Principal Component Analysis

PCA is a dimensionality reduction procedure which produces a series of orthogonal reaction coordinates from linear combinations of the input data *x*_*i*_, *i* = 1, 63. Its aim is to retain and explain as much of the variance of the original data as possible. This procedure, first introduced in^34,35^, relies on the solution of an eigenvalue problem involving the covariance matrix of the input data

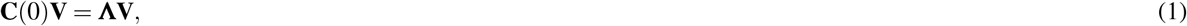

where **C**(0) is the covariance matrix *C*(0)_*ij*_ ∝ ∑_*t*_ *x*_*i*_(*t*)*x* _*j*_(*t*) of the normalized data (mean 0, unit variance), **V** contains the eigenvectors *v*_*i*_ by columns, and **Λ** is the diagonal eigenvalue matrix. The *principal components* are therefore obtained by projecting our original data into the different eigenvectors. A more detailed look into the derivation of the method can be found in the Supplementary Material.

The relative magnitude of the resulting eigenvalues is a measure of the proportion in which the original variance is projected onto the corresponding PC. Thus, depending on the distribution of the eigenvalues *λ*_*i*_, a subset of coordinates *n* ≤ *N* can be selected for the ongoing system description. The appropriate value for *n* corresponds to a large reduction in the magnitude of the eigenvalue *λ*_*n*+1_ when compared to *λ*_*n*_, meaning that most of the relevant information of the system (which corresponds to the majority of the variance) resides within the first *n* degrees of freedom.

In our specific context, PCA is employed to eliminate extraneous degrees of freedom while retaining those that contribute to insights into conformational changes within the G4 structure. It is crucial to note that this assumption holds true only when these transitions represent the most significant alterations in the variance of the system, a condition that may not always be met. To selectively filter changes based on their kinetics and retain only those with a timescale exceeding a predefined threshold, we employ tICA, here below presented.

#### 2.3.2 time-Independent Component Analysis

tICA is a method, first introduced in^36^, which produces a series of reaction coordinates from linear combinations of the input data that maximise not the variance, but the autocorrelation of the projected data between times t and t + *τ*, with *τ* being a time window selected by the user, known as the *lag time*. The mathematical framework of tICA is similar in its nature to that one of PCA, and is presented in the Supplementary Material.

The method relies on the resolution of a generalized eigenvalue problem involving the covariance matrix of the data and the time-lagged correlation matrix *C*(*τ*)_*ij*_ ∝ ∑_*t*_ *x*_*i*_(*t*)*x* _*j*_(*t* + *τ*), similar to the former but correlating the data at times t and t + *τ*:

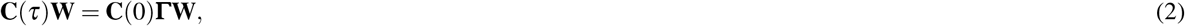

where **Γ** is the diagonal eigenvalue matrix, and **W** the eigenvectors matrix, with each of the different *w*_*i*_ vectors as columns. This general equation (Eq. (2)) is typically not solvable due to the small value of the determinant of the matrices involved, leading to numerical errors in the calculations. The AMUSE algorithm^59^ is typically used in its place, detailed in the Supplementary Material.

It has been shown^60^ that the eigenvalues *γ*_*i*_ correspond to the value of autocorrelation of the *i*-th component and that the cross-correlation between the *i*-th and the *j*-th coordinates vanishes at the lag time *t* = *τ*. Furthermore, if we assume the autocorrelation of the signal to have an exponential decay, the associated constant is given by the lag time and the eigenvalue as:

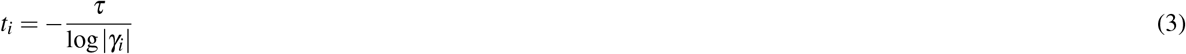

Therefore, the value of the lag-time acts as a threshold for the time-scales detected in our system: if a certain timescale is bigger than *τ* it will be detected and included in our coordinates, otherwise it will be discarded, since its autocorrelation has had time to decay to 0: it is a kinetic filter. With this property in mind, tICA should be able to distinguish important conformational changes and intermediates that PCA could not consider if they are not affected by a remarkable variance value.

### 2.4 Conformational Markov Network and conformational basins

Conformational Markov Networks (CMNs) are a tool that has been previously introduced to study the Free Energy Landscape of a diverse array of physical systems^37–40^. It consists in building a series of nodes connected via directed links.

For the construction of a CMN, each of the *n* reduced coordinates, obtained from either PCA or tICA, are discretized into *m* intervals. Every possible combination of the intervals of the coordinates that has been occupied (i.e. the trajectory of our real system has been in that precise combination at an arbitrary time *t*) will constitute a *node* in our system. Therefore, we have a number of nodes *N*_nodes_ ≤ *m*^*n*^. The weight of each node is given by the number of times its associated combination of intervals is visited by the trajectory, normalized with the total number of time frames the trajectory is divided in. The links between them are directed and described by *P*_ij_, denoting the transition probability from node *j* to node *i*, normalized such that ∑_*i*_ *P*_ij_ = 1.

The resulting network size can become challenging to analyze depending on the chosen number of intervals *m*, which is directly correlated to the number of nodes. In our specific case, *n* = 3 coordinates and *m* = 10 intervals were chosen, resulting in up to 1000 distinct nodes. To reduce this number and eliminate potential redundancies where different nodes essentially represent the same conformation, a Stochastic Steepest Descent algorithm was employed. This algorithm, described in detail in Ref^37^, groups the nodes into basins, which represent different attractors of the trajectory. Additionally, a filtering based on a cutoff of the weight of the nodes has been introduced with the goal to retain only fundamental basins, and eliminating the ones less representative (*P*_*i*_ *<* 10^−5^).

## 3 Results

### 3.1 Direct GROMACS simulations

One of the magnitudes we investigated in our study is the *Root Mean Square Deviation* (RMSD) of the G-quadruplex structure: the difference at each time between the G4 positions and the equivalent native configuration, calculated along all replica’s trajectories. This measure is the simplest way to indicate how much the structure changes during the time evolution, and it has been calculated both for the complete G4-system and the guanine tetrads only (*i*.*e*. the piled guanine structure without the external loops), the latter being the most important guideline to confirm that changes in the RMSD actually correspond to unfolding processes. In fact, a significant change in loop conformation could lead to an increase in the total RMSD, while not necessarily being associated to the denaturation of the G-quadruplex. Thus the confirmation of the unfolding relies on the G-tetrads structure only.

This information is contained in Fig. 2, that shows both the whole G-quadruplex and G-tetrads. To display the RMSD values properly, we have *d*emultiplexed the trajectories: instead of tracking the behavior of a replica whose contents are continuously changing over time, we analyze the evolution of a specific G4 structure as it passes through between replicas. This approach prevents the misinterpretation of changes in RMSD that are actually due to exchanges between replicas as real unfolding events. We refer to these *d*emuxed trajectories as *Systems* (Sys.), as they represent the same nucleic structure over all the replicas. The successful exchanges between replicas, when a Sys. changes temperature, are marked by gray vertical lines in Fig. 2. The RMSD curves focusing on the replicas can be found in the Supplementary Material, corresponding to Supplementary Figs. 1 & 2.

**Figure 2.**
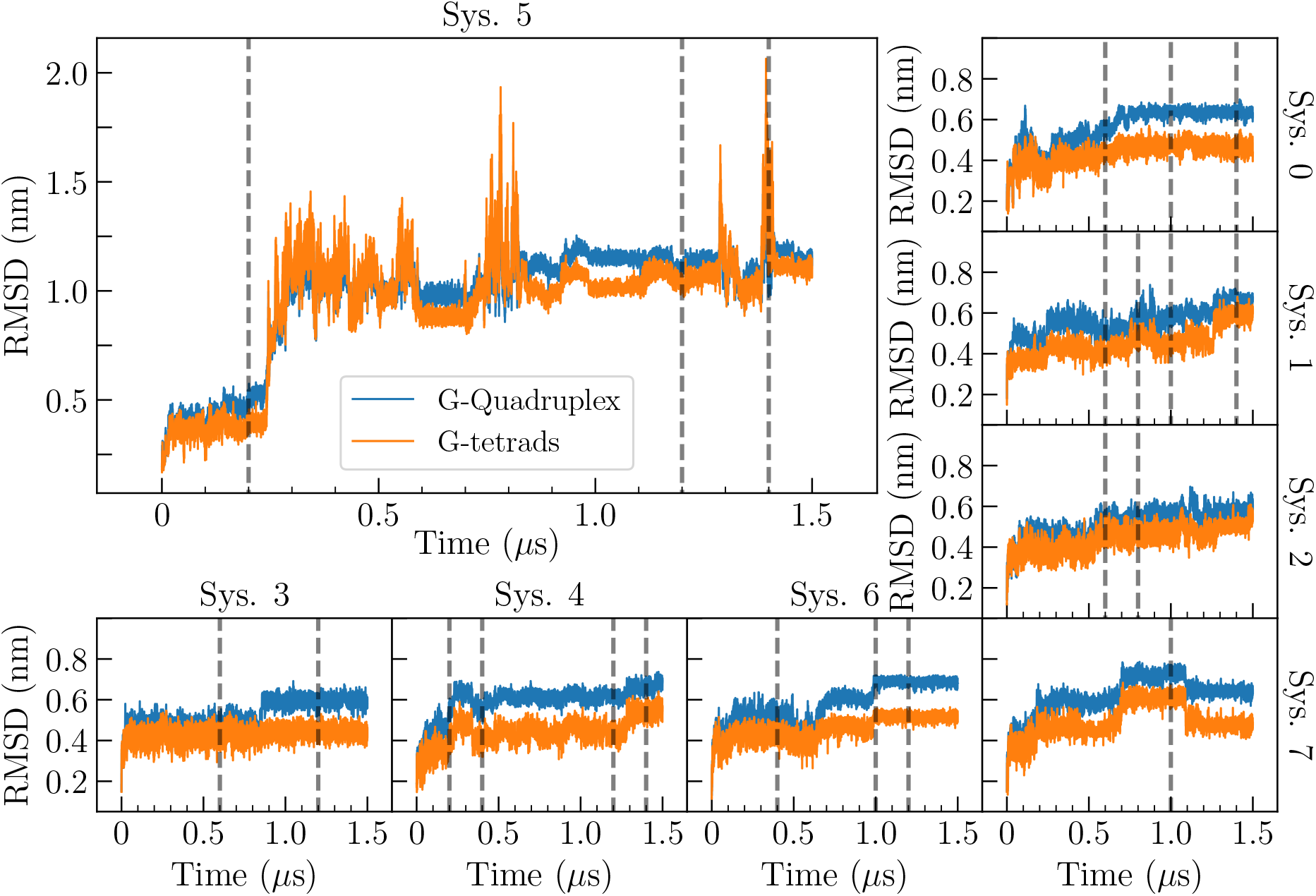
RMSD calculated over the different starting configurations (Sys.) of the parallel G-quadruplexes. In blue, the RMSD of the whole structure. In orange, the RMSD taking into account only the guanines forming the planar arrangement. The vertical dashed lines correspond to successful exchanges between replicas. Sys. 5, since its the one experiencing unfolding and used for the analysis, has been enhanced.

All but one of the Sys. present an RMSD that quickly stabilizes at either approximately 0.6 nm or 0.4 nm, depending either on the inclusion (complete G4, blue line in the panels) or absence (G-tetrads, orange line) of the loops in the calculation, respectively. Sys. 5 undergoes great deviations from the norm, reaching values up to 2 nm independently of the loops, thus confirming an unfolding event. These results further emphasize the correctness of the choice of the REMD method; in fact, the unfolding processes of the G-quadruplex have an average lifetime typically larger than the scope of the simulations, thus making them difficult to observe without the use of replicas at different temperatures.

The other G4 structures remain in a relatively stable configuration, though some of them present the loss of one ion from the central channel, occurrence described in the next Section 3.1.1.

These same results can be understood through a second metric, the *radius of gyration* (R_g_), which measures the average square distance of the monomers respect to the center of mass of the structure. Fig. 3 shows the values of (R_g_), as a function of time, where we can observe its low variability in the majority of systems, while a rapid and sudden deviation with respect to the stable value occurs in Sys. 5, indicating the unfolding of the structure already seen in Fig. 2.

**Figure 3.**
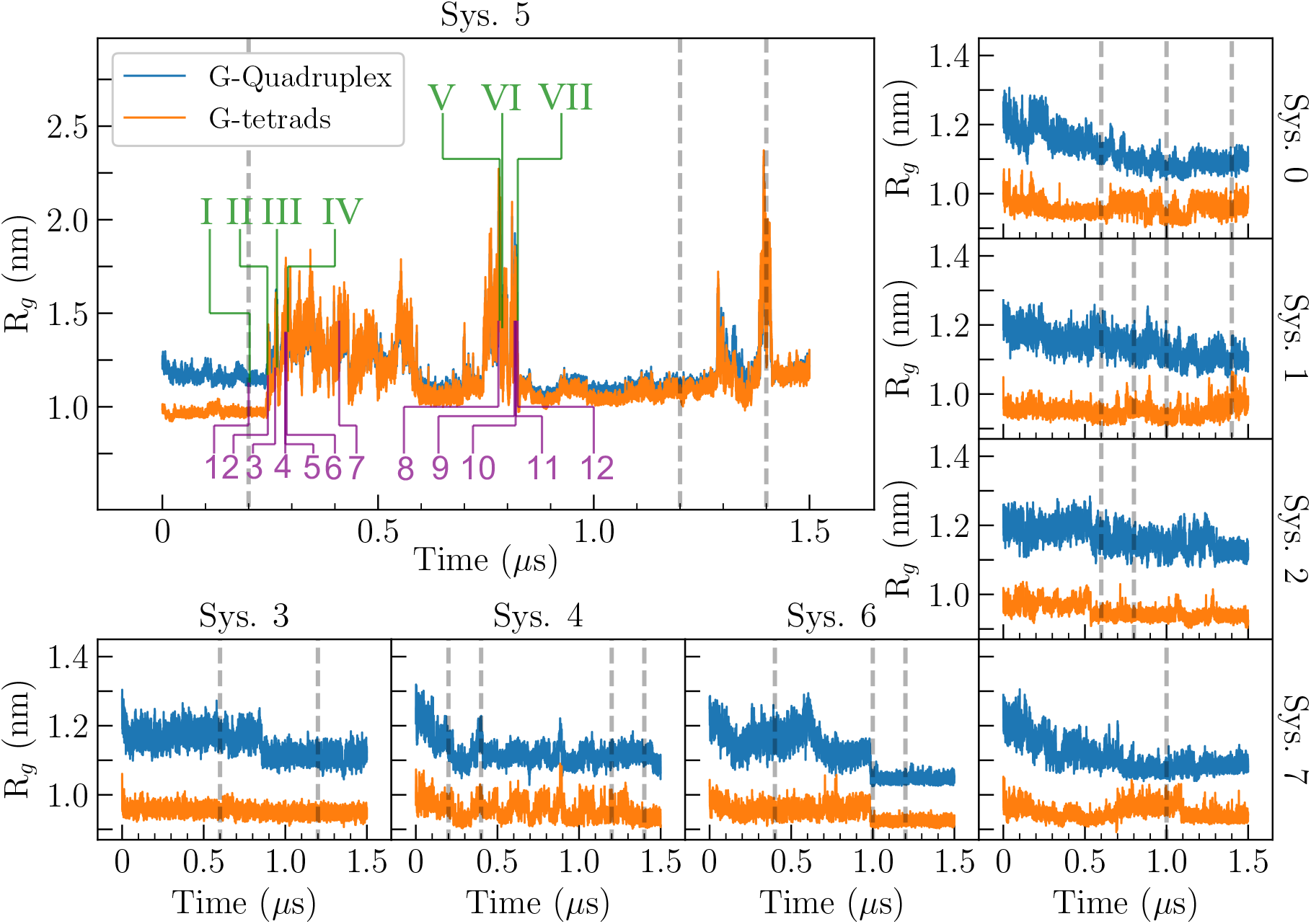
Radius of gyration computed over the different starting configurations (Sys.) of the parallel G-quadruplex. In blue, the R_g_ of the whole structure. In orange, the R_g_ for the G-tetrads. Sys. 5 has been enhanced, since it is the one considered for posterior analysis due to the presence of unfolding. The vertical grey dashed lines correspond to successful exchanges between replicas. The purple and green colors of the numbers correspond to the states identified by PCA and tICA respectively, corresponding to the conformations reported in Figs. 10 & 11.

#### 3.1.1 Effect of the monovalent ions in the stability of the system

The stability of G-quadruplex structures is highly influenced by the presence or absence of ions, typically monovalent, within the central channel^6^. These ions, which stably reside in the G4 because of their positive electrostatic interactions with the negatively charged G-tetrads and the spatial arrangement of guanines in the native conformation, may escape if a structural conformation with a sufficiently wide central channel occurs.

The replica analysis is once again inverted, focusing on a single structure as it traverses between replicas to prevent confusing exchanges between replicas with ion loss events. To characterize the ions’ position, two coordination numbers were employed: one for the site corresponding to the top and intermediate planes, denoted as Co_1_, and the other for that of the intermediate and bottom planes, denoted as Co_2_. These metrics quantify the number of bonds formed between each ion (filtered according to the closeness to the G4 structure) and the guanine bases constituting the tetrads, with values ranging from 0.0 (indicating no bonds) to 1.0 (indicating all 8 possible bonds) in increments of 0.125. A bond is considered established if the distance between the O6^*′*^ atom of guanine and the ion is less than 4.5 Å.

Since the middle tetrad is involved in the calculation of both coordination numbers, an ion exhibiting a coordination number of 1.0 in one of the two possible placements would also present a value of 0.5 in the other. A situation like this could be mistaken with the presence of two ions within the structure, one fully coordinated (hence Co_i_ = 1.0) and another partially coordinated (Co_j_ ≤ 1.0, *j* ≠ *i*). To prevent this specific confusion, once an ion achieves Co_i_ = 1.0 in one position, its coordination number is set to 0 in the other position. Monitoring the width of the central channel itself involves defining an additional set of coordination numbers for the three guanine tetrads, referred to as Co_P_. These numbers, similar to those used for ions, quantify the distances between the four guanines forming each tetrad, based on the Hoogsteen bonds present in the structure. The coordination number ranges from 0.0 to 1.0 in increments of 0.25, with a bond considered established if the distance between two neighboring guanines is less than 5.0 Å.

Fig. 4 illustrates the evolution of the two coordination numbers for the ions (Co_1_ and Co_2_ in the first two rows), along with the three coordination numbers of the G-tetrads (third row, each color represents a plane as depicted in Fig. 1: blue for the top plane, red for the middle plane and green for the bottom plane) and the radius of gyration of the G-tetrads (loops excluded). All systems, except Sys. 3, experience the loss of one or both ions at some point of their trajectories. Excluding Sys. 5, which unfolds and consequently lacks a definable central channel, the systems either recover both ions or maintain one ion while losing the other, as observed in Systems 0, 1, 2, 4 and 7. It should be noted that the recovered ions are not necessarily those initially lost; in fact all the ions in the simulation box are equivalent, thus color changes in Fig. 4 can eventually occur. The departure of an ion from the channel increases the system’s susceptibility to destabilization. This phenomenon is reflected in Fig. 4, where the loss of an ion from either position in the channel leads to significant fluctuations in both the coordination number of the corresponding planes and the radius of gyration R_g_. An example of this can be seen in Sys. 7 in the time window [0.75 - 1.0] *µ*s, showing fluctuation in the plane coordination numbers and deviations in R_g_. The same deviations can be seen in Rep. 7 in Figs. 2 & 3 during the same time interval.

**Figure 4.**
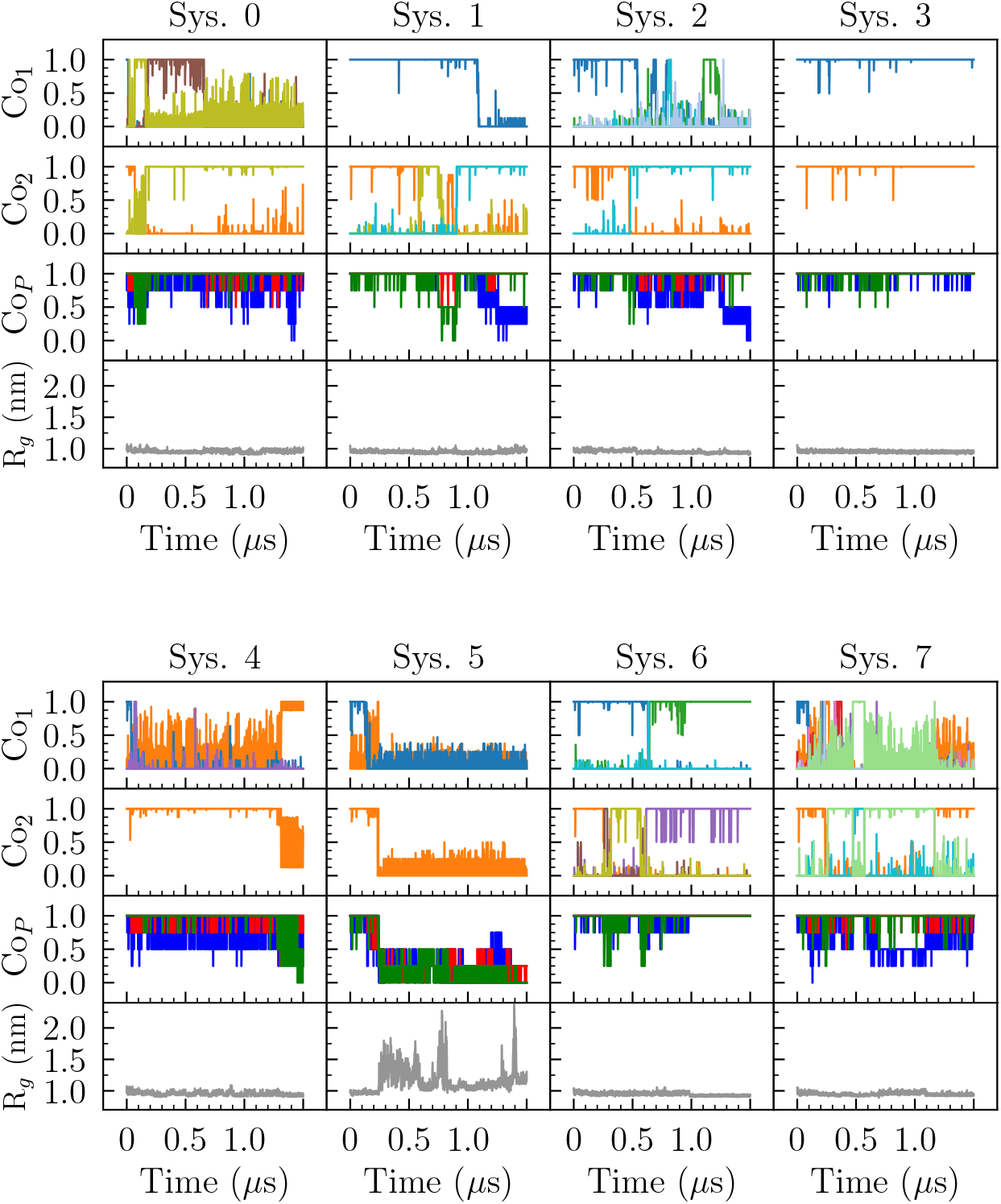
Coordination numbers of the top (Co_1_) and bottom (Co_2_) positions available for the ions, coordination number of the three G-tetrads (Co_P_, blue for the top plane, red for the middle plane and green for the bottom plane) and radius of gyration of the G-tetrads (loops excluded). Only ions which achieve a value of 1.0 in each system are represented.

Ion escape and reabsorption are facilitated by slight changes in the central channel’s width. The primary causes of these deformations includes the bending of G-tetrads, the motion of single guanines unbinding/drifting away from the structure. These deformations are not limited to the top and bottom tetrads. Disruptions in the intermediate plane can also result in the transfer of an ion from one position in the ionic channel to another, as observed in Sys. 0 at 0.2 *µ*s, Sys. 4 at 1.3 *µ*s and Sys. 7 at 0.5 *µ*s.

Fig. 5 shows various conformations associated with the dynamics of the ions in the different systems. Subfigure **A** illustrates the escape of an ion from the top position, corresponding in Fig. 4 to the fall of the blue line in Sys. 1 around 1.1 *µ*s, which triggers a decrease in Co_P1_. Subfigure **B** shows the reverse process, where an ion is integrated into the structure, associated with the increase of the olive Co_1_ line at around 0.2 *µ*s in Sys. 0 and a corresponding rise in Co_P2_. Subfigures **C** and **D**, taken from Sys. 2 and 7, respectively, depict the separation of a guanine from the top (blue) plane after ion escape around 1.3 *µ*s and 0.7 *µ*s respectively, leading to a significant drop in the coordination number of that plane, Co_P1_. Subfigure **E** shows a similar phenomenon, but with the displaced guanine in the bottom plane. This separation occurs while an ion remains bound, eventually leading to its loss around 0.76 *µ*s in Sys. 2, accompanied by a decrease in Co_P3_. Lastly, Subfigure **F** shows a slip-stranded conformation in Sys. 4 at 1.3 *µ*s. The slip-stranding effect reduces the maximum possible coordination number to 0.875 and widens the central channel, facilitating the transfer of the ion from the bottom to the top position, altering the trends of the coordination numbers. This slip-stranding also causes a decrease in the coordination number Co_P3_, with a new maximum value of Co_P3, max_ = 0.75.

**Figure 5.**
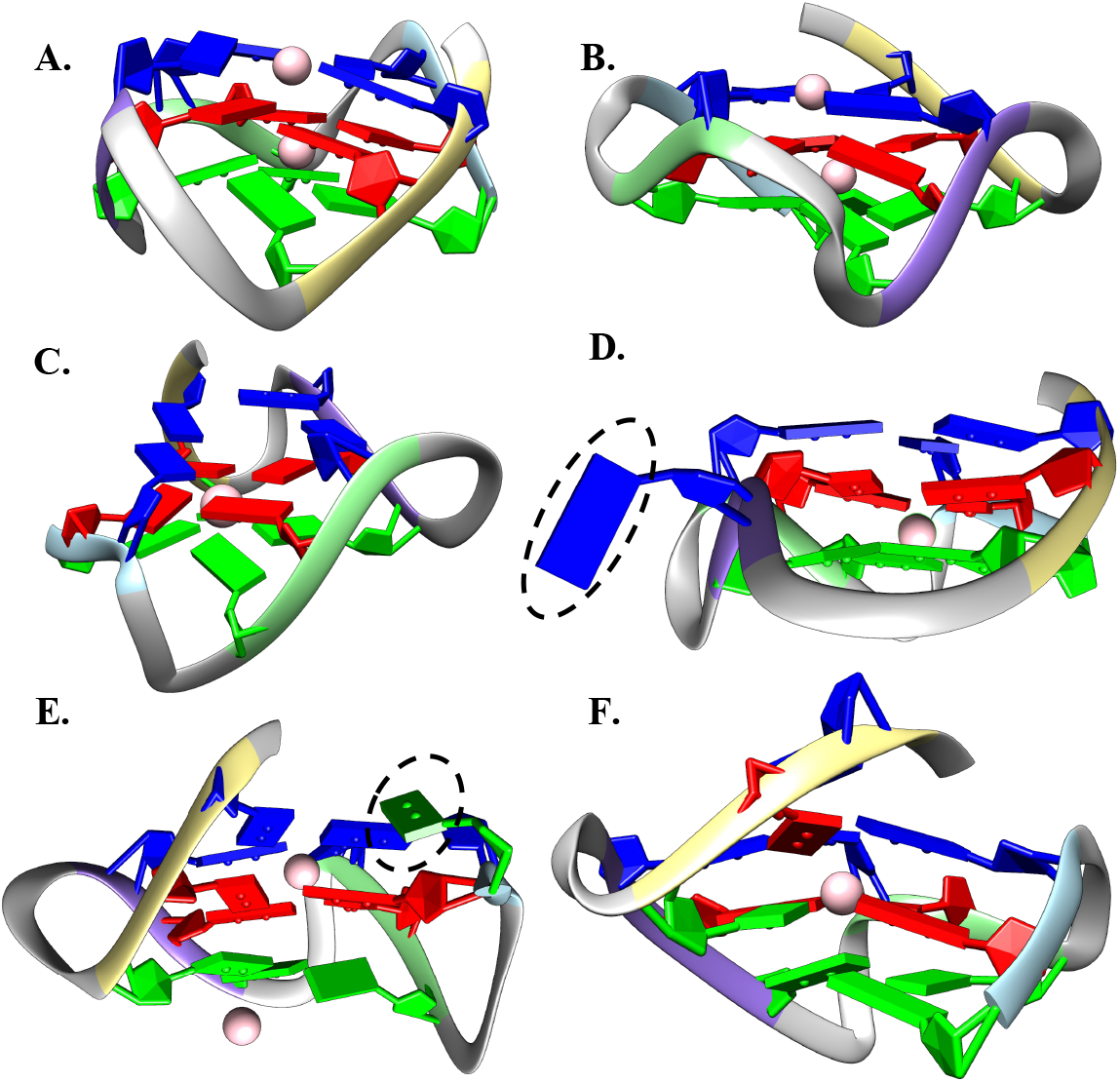
Relevant conformations associated to ion dynamics in the simulation. **A.** Escape of an ion from the top plane. **B**. Absorption of an ion through the top plane. **C**. Deformation of the top plane after ion loss. **D**. Separation of a single guanine from the tetrad of the top plane (indicated by black dashed outline). One ion has already left the structure. **E**. Separation of a single guanine from the bottom tetrad (indicated by black dashed outline), leading to ion loss. **F**. Slip-stranded conformation leading to the transference of an ion from the bottom to the top parts of the central channel. Slip-stranded contacts circled by dashed lines.

From these results, it is evident that the presence of ions significantly influences the G-quadruplex stability, generally leading to partial guanine separation and other structural changes following their loss. In Sys. 5, which undergoes unfolding, both ions are lost before the process begins: one is lost early in the simulation, briefly replaced by another, which is lost again around 0.1 *µ*s, and the other ion is lost at 0.24 *µ*s, with unfolding commencing around 0.26 *µ*s.

In the remaining systems, ion loss typically occurs in only one of the two available cavities. In this sense the loss of only one ion make the G4 structure less stable as it appears like a metastable state that maintains a certain stability. The G4 structure remains not completely compact, as visible in Fig. 5, where the structures are partially modified with the loss of one ion. We understand that these structures can maintain relative stability over time up to the loss of the second ion, a condition that definitely triggers the unfolding process. This observation may explain why only one system fully unfolded, as it required the loss of both ions before the unfolding process could initiate.

#### 3.1.2 Unfolding process

The unfolding process occurs in Sys. 5, shown in Figs. 2 and 3. The identification of peculiar states in the trajectory will be largely developed with the analysis with tICA and PCA, later on presented.

In Sys. 5, the initial configuration after the replica exchange occuring at *t* = 0.2 *µ*s corresponds to a folded G4 which has already lost one of its ions, as visible in Fig. <u>4</u>. Afterwards, it quickly loses the other ion, at *t* = 0.24 *µ*s, therefore triggering the unfolding itself, as reflected by the increase of RMSD and R_g_ in Figs. 2 & 3 at *t* = 0.25 *µ*s.

Fig. 6*α* presents a G4 conformation in which the fourth guanine tract (ice blue, see Fig. 1) separates from the structure with the remaining three segments, thus forming a G-triplex^61,62^, one common intermediates in G-quadruplex unfolding^63,64^. During the next 30 ns several attempts at refolding are made unsuccessfully. Moreover the dashed circle in Fig. 6*β* shows, in the remaining G-triplex, the so-called “slip-stranding”, *i.e*. a guanine tract that moves upwards or downwards respect to the others^65,66^. Other transient conformations can appear, such as G-hairpins^63,67–69^, i.e. the bond of two guanine tracts only (see Fig. <u>6</u>*γ*), or even temporary refolding events into a G-triplex (Fig. 6*δ*).

**Figure 6.**
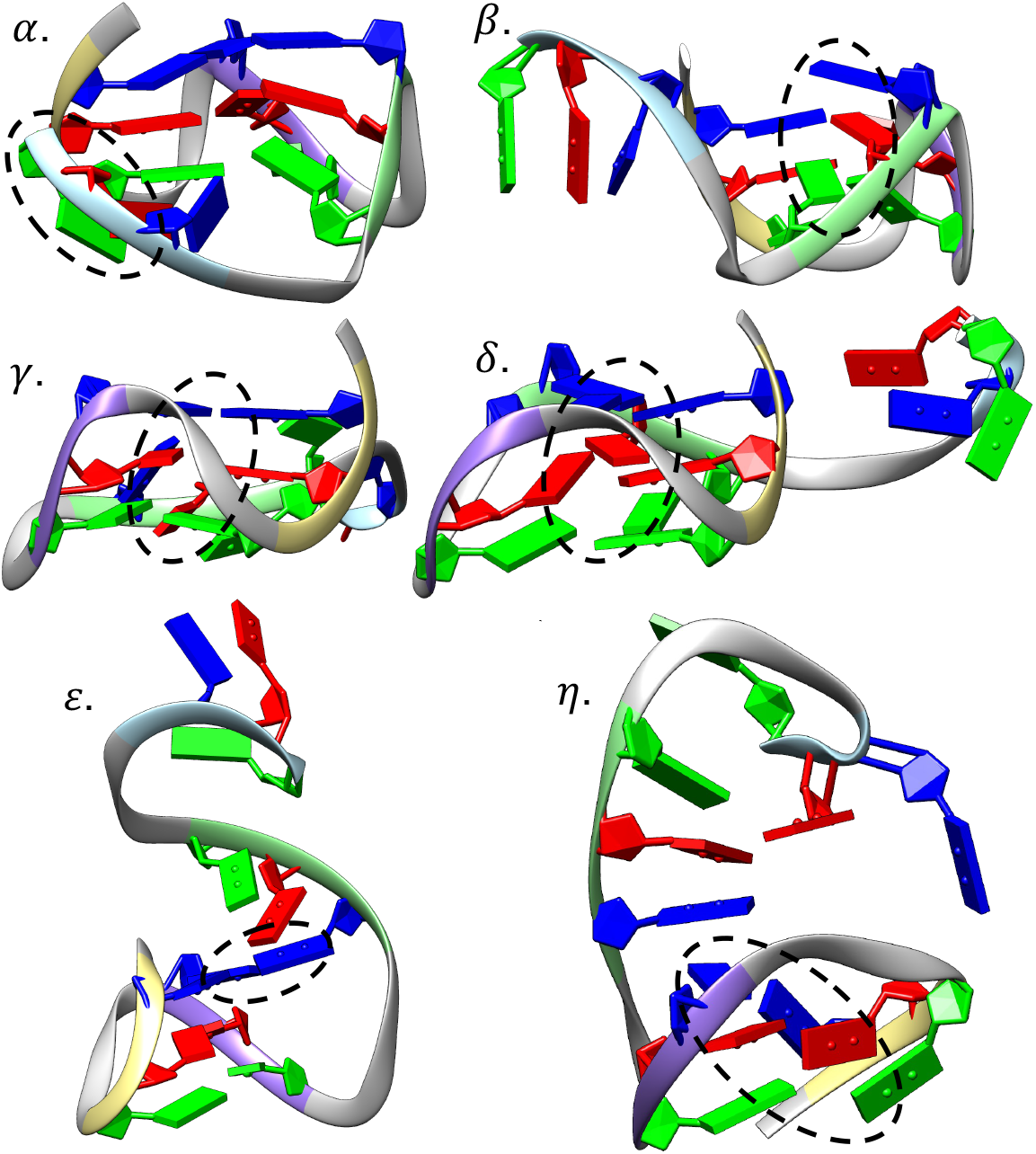
Highlighted conformations achieved during the evolution of the system, with the dotted circles highlighting the features of interest. ***α*.** Initial detachment of the fourth (icy blue) tract of guanines. ***β***. Example of a G-triplex conformation with a slip-stranded effect between tracts 1 (yellow) and 3 (green). ***γ***. Formation of a G-hairpin between tracts 1 (yellow) and 2 (magenta). ***δ***. Refolded triplex after re-attachment of tract 1. ***ε***. Stable intermediate state of G-hairpin (tracts 1 and 2) with stacking interaction of the third tract. ***η***. Final state of the system, with tracts 1 and 2 in a cross hairpin conformation.

However, after the successful refolding into a triplex, the third G4 tract detaches again, rotates over itself and forms a single Hoogsteen bond with the top plane of the G-hairpin formed by the first and second tract, as shown in Fig. 6*ε*. Remarkably, this latter hybrid between a G-triplex and a G-hairpin remains stable for approximately Δ*t* ≈ 0.6 *µ*s of the simulation. The intense fluctuations observed in Fig. 3 in the interval [0.3, 0.9] *µ*s correspond to changes in the distance between the third and fourth tracts. Eventually, the fourth strand rotates over itself and binds with the third strand through the bottom guanine (green), corresponding to the plateau between *t* = 0.6 *µ*s and *t* = 0.75 *µ*s. After instant *t* = 0.75 *µ*s, the bond joining the G-hairpin with the third tract breaks. The chain elongates, pulling in the second tract and separating it from the first, breaking the G-hairpin. It is quickly reformed but with the two tracts rotated, forming a “cross-hairpin”^69^. During this time, the third and fourth tracts rearrange themselves into two main conformations: one characterized by the stacking interactions between both tracts leading to a high R_g_, and another presenting the formation of Hoogsteen bonds between their guanines, with smaller R_g_. The interchange between them forms the peaks observed around 0.8 *µ*s.

The final conformation of the system, stable and leading to the plateau in Fig. 3 starting at *t* = 0.8 *µ*s, depicts the formation of a *cross hairpin* (an arrangement of guanine rich strands in a cruciform shape, as circled in Fig. 6*η*) between the first and the second tract, with the third and fourth stacks remaining close, accompanied by the formation of transient Hoogsteen bonds between them.

### 3.2 Dimensionality reduction

Both PCA and tICA are techniques quite sensible to the characteristics of the data of interest.

The application of the two methods in our trajectories has required to clean the data as follows: i. first of all, we corrected the effects of the periodic boundary conditions on the GROMACS trajectories by removing the artificial discontinuities; ii. in order to prevent unrealistic distances of the trajectory coordinates emerging from global rotations and/or displacements of the structure as a whole, the coordinates of the structure have been rescaled by applying both a translation, superimposing the centers of mass with that of native structure, and rotations of the whole structure to recover the native orientation. A mean-square distance method, taking the initial state of the system as a reference, has been applied. iii. Finally, the mean was subtracted from each of the trajectory coordinates *x*_*i*_, which have been used as input values for both PCA and tICA analysis.

#### 3.2.1 Eigenvalues

The eigenvalues obtained from solving Eqs. (1) and (2) give the possibility to reduce the information provided by the complete degrees of freedom of the system into a smaller set of variables containing its most significant part.

In PCA, the relative magnitude of the eigenvalues (with respect to their total sum) is related to the percentage of the total variance projected onto the corresponding eigenvector which is combination of the initial coordinates. Thus, the larger the eigenvalues, the more relevant that particular direction is to represent the overall description of the dynamics. Ideally, one should find an eigenvalue or a small series of them with magnitudes clearly superior to the rest, indicating that the majority of information of the variance of the system is contained in one reaction coordinate. The outcomes of the diagonalization of the correlation matrix are contained in the left side of Fig. 7.

**Figure 7.**
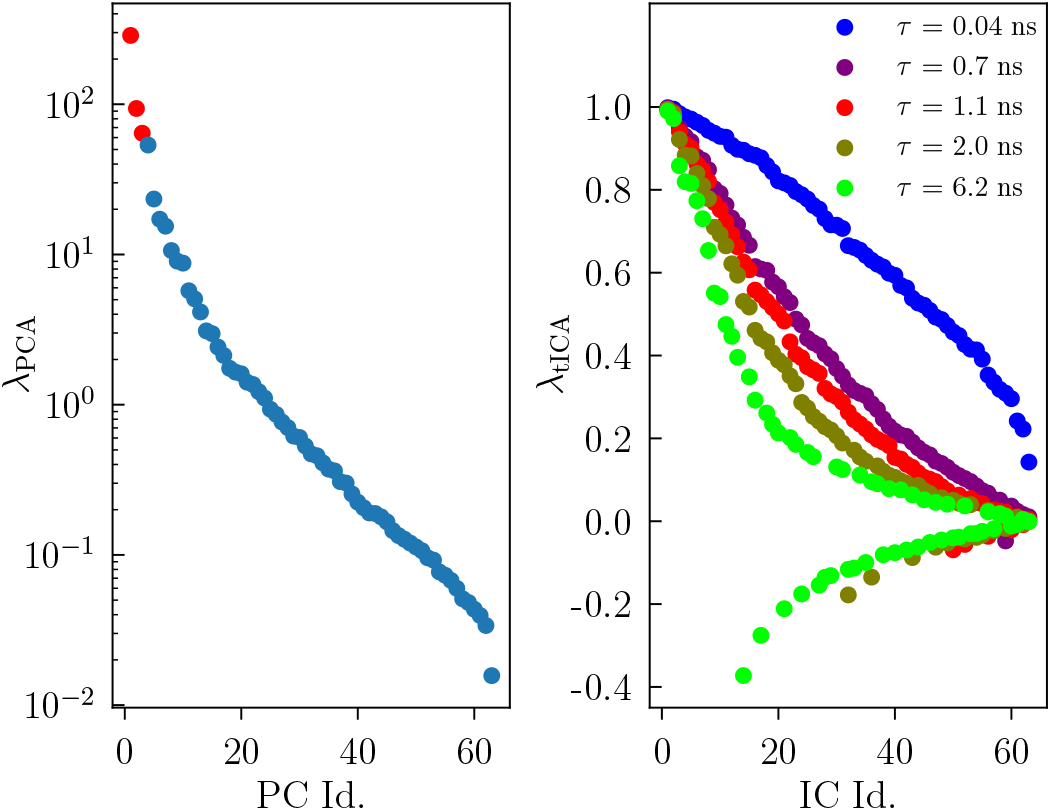
On the left, PCA eigenvalues obtained from solving (<u>1</u>), with the vertical axis in log-scale. The three eigenvalues in red correspond to the combinations chosen for data projection. On the right, tICA eigenvalues obtained from solving (<u>2</u>) for different values of the time window *τ*, seen in the legend.

The first four eigenvalues are clearly larger than all the rest, specially the first. We choose to use the first three eigenvectors associated to them as the basis for the PCA procedure, accounting for 70.9% of the variance.

For tICA, the interpretation of the eigenvalues is not as straightforward; their magnitude indicates the minimum timescale its associated coordinate is able to discern, calculated as in Eq. (3).

The eigenvalues of tICA are constrained between -1 and 1, with the negative values only appearing for large values of the lag time *τ* (see *τ* = 2.04 ns in Fig. 7). These eigenvalues indicate modes in the system (described by their corresponding eigenvectors) that decay over time, typically corresponding to fast transitions or back-and-forth fluctuations that do not contribute to the stable, meaningful changes tICA aims to capture. Thus, the appearance of negative eigenvalues at a certain lag-time *τ* imposes an effective ceiling onto this magnitude, *τ* = 1.1 ns in our particular case. The right side of Fig. 7 contains the eigenvalues for tICA calculated at different lag-times, showing the appearance of negative eigenvalues from *τ* = 1.1 ns onwards.

The analysis of the trajectories was carried out with different values of *τ* but, for the rest of this document, a lag-time of *τ* = 0.7 ns has been chosen. The reason for this particular value relies on the compromise we found which consists, on the one hand, in eliminating as many fast irrelevant motions of the system as possible that are translated into a few simulation instants and thus undetected by higher values of tICA, while, and on the other hand, in protecting the information contained in the longer-lived states.

#### 3.2.2 Projected trajectories

Fig. 8 shows the trajectories of both methods, PCA and tICA, projected on the eigenvectors associated to the three chosen eigenvalues with, on the right hand side, the histogram of the corresponding coordinates occupation. The analysis of Fig. 8 reveals appreciable differences between the two methodologies. PCA yields trajectories characterized by histograms exhibiting broad Gaussian-like peaks, whereas tICA produces trajectories with narrower distributions, able to better distinguish between different states of the system with improved clarity. Commonalities are observed in both methods, visible, for example in the first coordinate PC1 and IC1 which show a similar trend: high value for the first interval between *t* = 0.25 and *t* = 0.8 *µ*s, and after that a narrow transition to another constant value. Both trajectories describe the general behavior observed in the root-mean-square deviation (RMSD) (see Fig. 2), with tICA exhibiting less fluctuations than PCA, that, instead, almost reproduces the same shape as the original trajectory.

**Figure 8.**
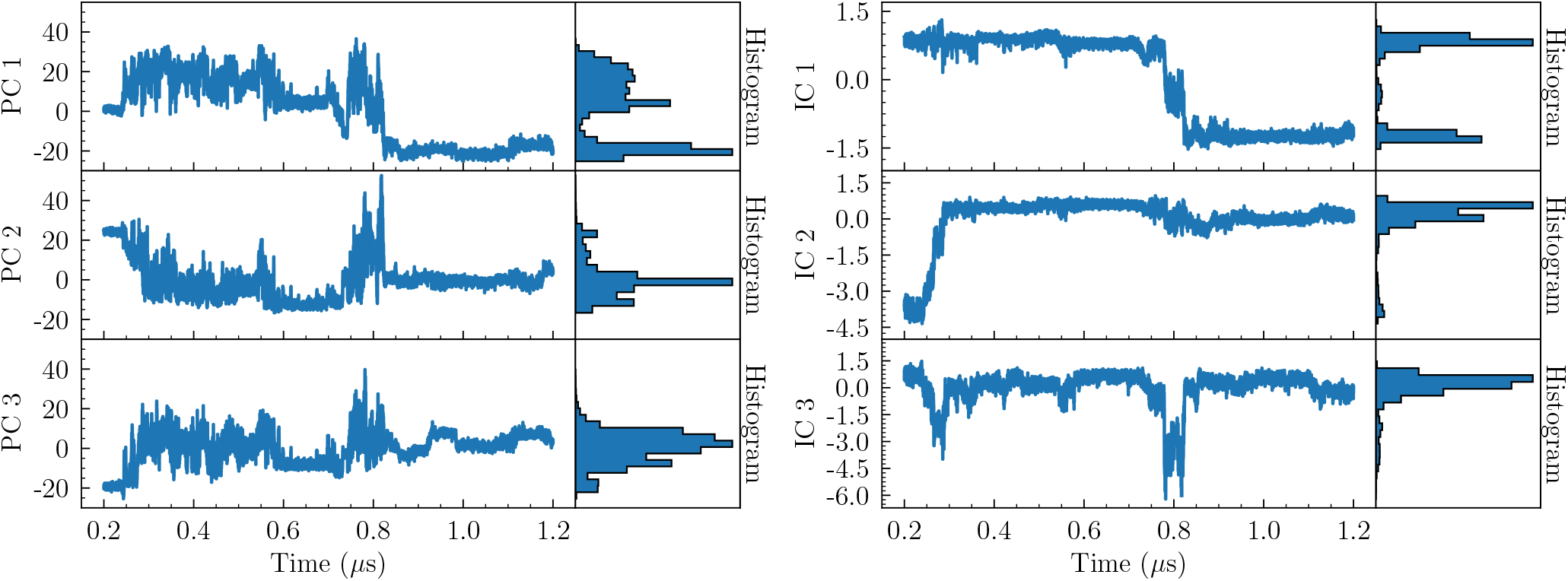
Projected trajectory from Sys. 5 between 0.2 and 1.2 *µ*s onto t he first three eigenvectors (left), along with the histogram of the coordinates (right). PCA on the right, tICA with *τ* = 0.7 ns on the left.

The different capabilities of the methods in detecting independent and well-defined states throughout the system’s trajectory become evident when plotting the coordinates against each other as in Fig. 9. The image depicts a 3D plot of the coordinates extracted from PCA (left) and tICA (right), with the color scale corresponding to the free energy differences in the trajectory determined by the relative occupation of specific coordinate combinations during evolution, and computed as − log(*P/P*_0_), with *P*_0_ being the lowest occupation probability (≠ 0) in the states ensemble. On the bottom of each 3D plot there are three heatmaps plotting the coordinates against each other by pairs.

**Figure 9.**
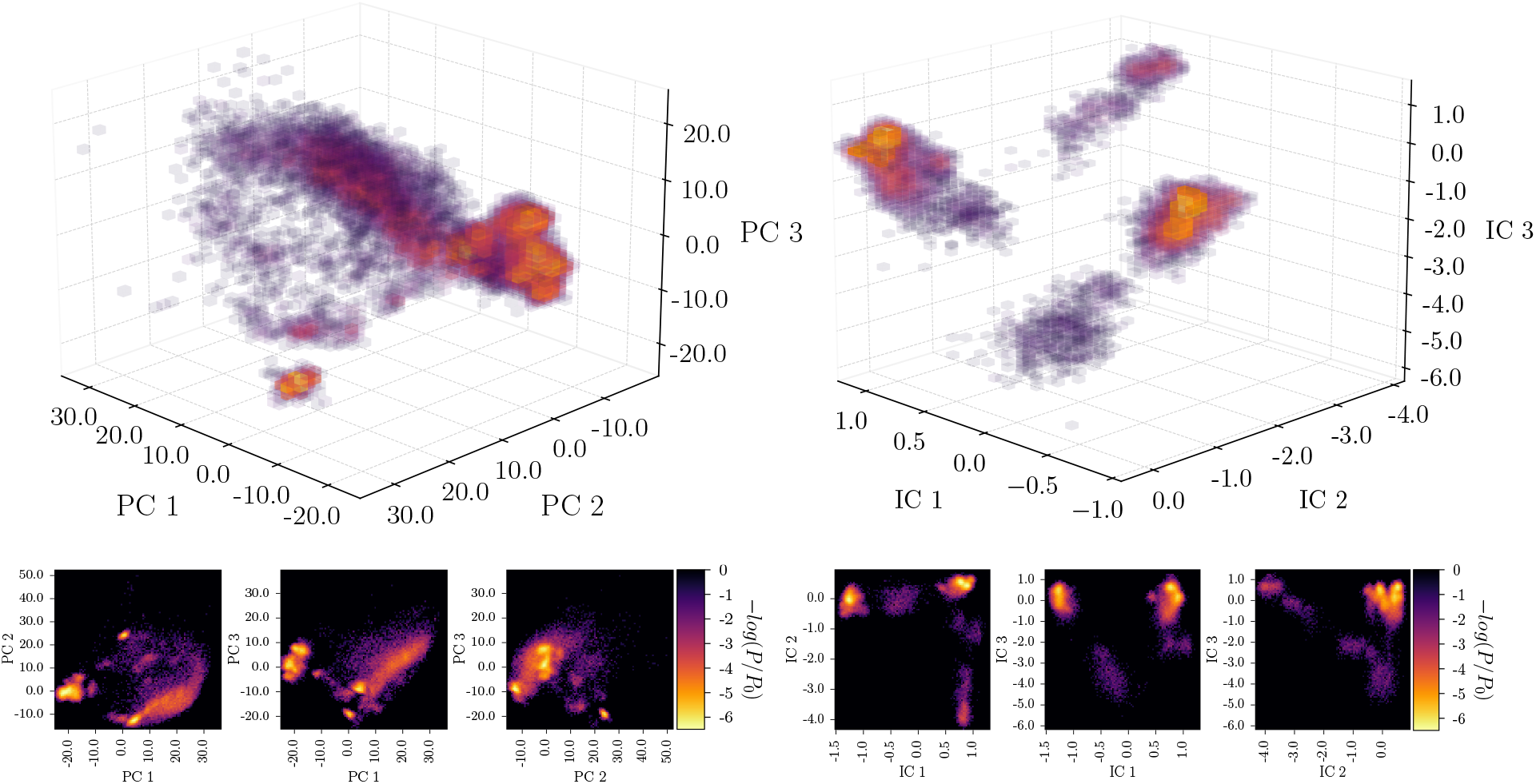
Colormaps depicting the correlation of the different coordinates emerging from PCA (left) and tICA (right). Within each subfigure, the bottom three panels depict heatmaps illustrating the pairwise relationships between the projected coordinates, with the color representing the relative occupancy of the corresponding coordinate pairs throughout the trajectory. The three-dimensional plots illustrates the combined coordination of the coordinates. Each data point in this plot corresponds to a specific conformation sampled during the trajectory, with the color representing the relative occupancy of a particular combination during the simulation.

In Fig. 9 is well visible that PCA exhibits greater variance in the evolution of its coordinates featuring bright states surrounded with clouds of relatively populated spots which, when 3D plotted, seem to collapse in a central globule. Conversely, tICA produces distinct bright spots that are well separated, with additional small areas of low population in between, leading to a 3D plot characterized by small cloud separated from one another.

### 3.3 Complex Markov Network results

Once a suitable value for *τ* for tICA is chosen and the trajectories are projected onto the reduced coordinate space we can proceed with their encoding into a complex network. As previously explained, the trajectory undergoes segmentation into equidistant subsets, each constituting a node when populated by the system. Subsequently, a Stochastic Steepest Descent algorithm is applied to facilitate the classification of all nodes into distinct basins of attraction.

The resulting networks of basins depicting the unfolding process are illustrated in Figs. 10 and 11.

**Figure 10.**
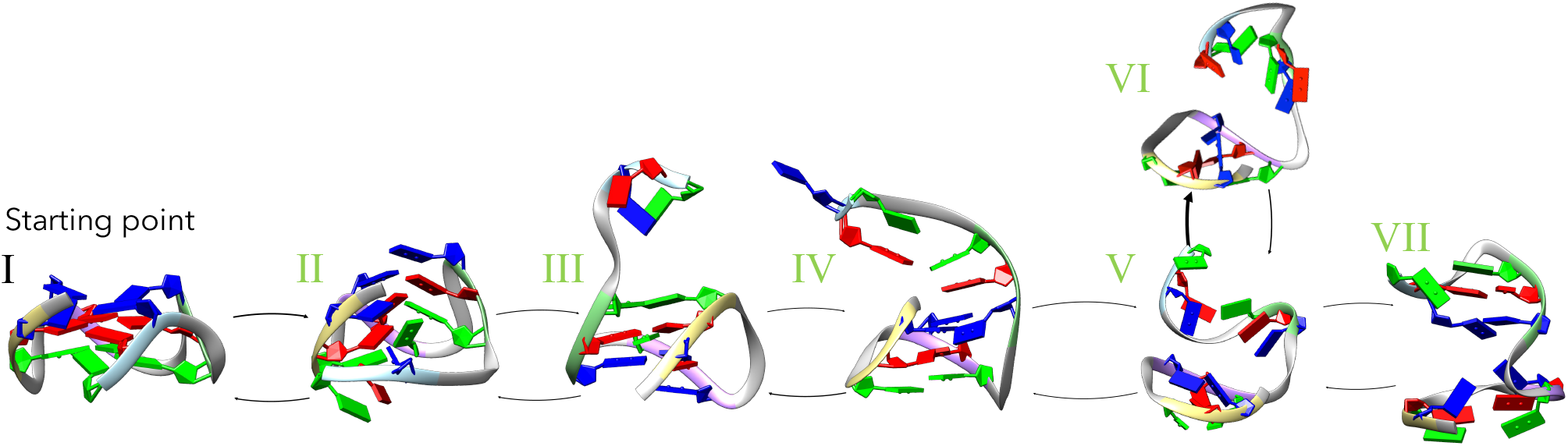
Network of basins found by tICA, *τ* = 0.7 ns. The green numbers refer to the equally colored lines in Fig. 3, showing the position of the basins along the trajectory. The files corresponding to the 7 structures, in.pdb format, as well as the videos of the unfolding, can be downloaded from the Supplementary Material.

**Figure 11.**
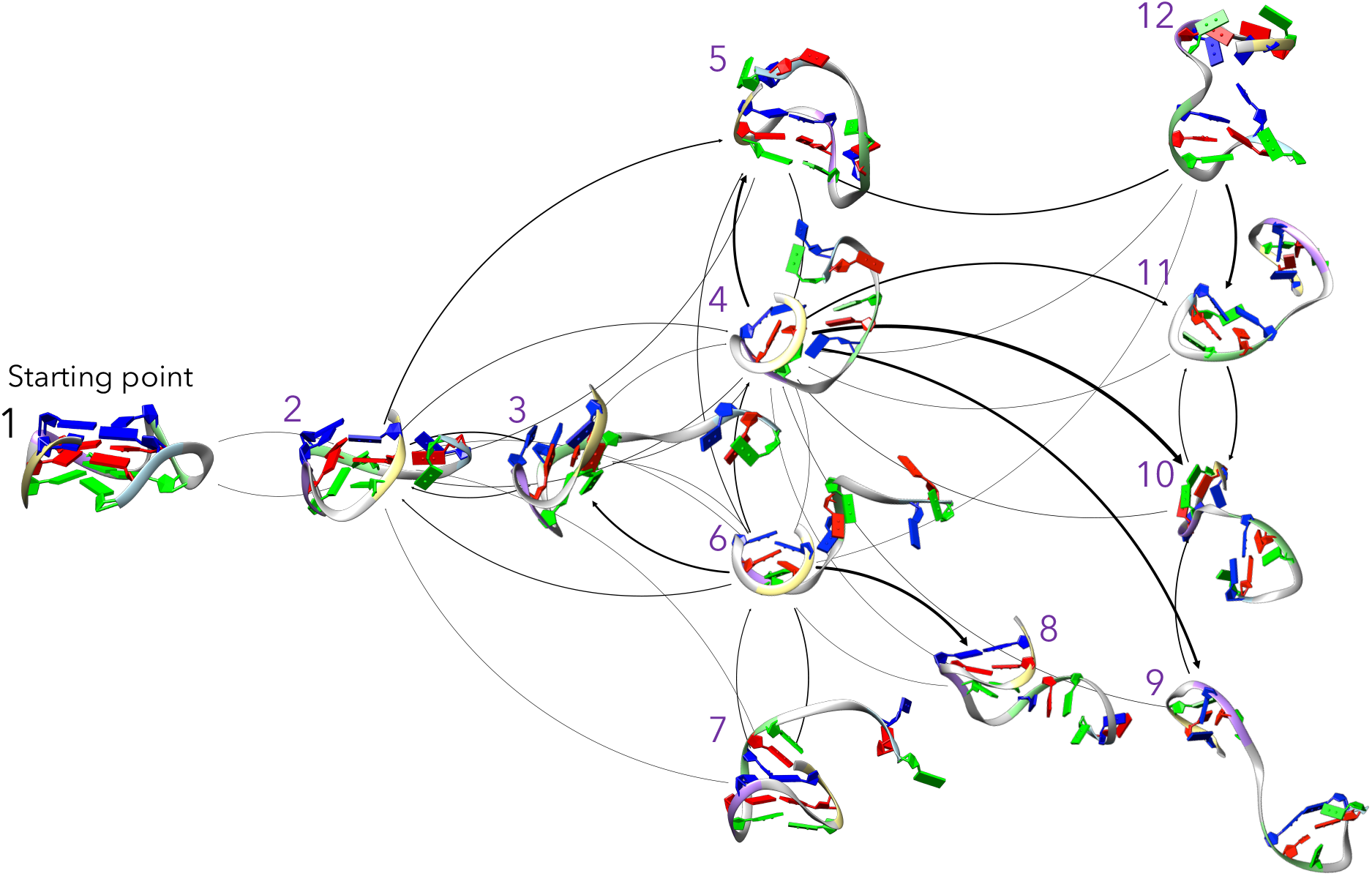
Network of basins found by PCA. The purple numbers refer to the equally colored lines of Fig. 3, indicating the position of the basins along the trajectory.

The results from tICA are presented first, since they provide a more straightforward depiction of the unfolding process. The PCA results are then analysed subsequently.

#### 3.3.1 tICA analysis

The tICA analysis summarized in Fig. 10 shows a linear representation of the unfolding process, with 7 states identified as basins of attraction in the CMN analysis. The representative node of each basin is labeled from 1 to 7 in the figure, and its corresponding time of appearance in the system trajectory is marked in Fig. 3 with green lines.

Starting from a fully folded conformation (basin #I), the next state (basin #II) portrays the system as already having severed its Hoogsteen bonds with the fourth guanine tract at the 3^*′*^-end of the chain. In this basin tICA is able to recognize the “slip-stranded” conformations between the remaining tracts in the G-triplex, reminiscing of Fig. 6*β*.

Basin #III depicts the moment after which the fourth guanine tract, previously disconnected from the main triplex but still proximal to it due to crowding effects, has separated. Moreover, another detail can be observed, consisting in the slip-stranding of tract 2 from the remaining triplex (slip-stranded contacts circled in dashed line in Fig. 10, basin #3).

In Basin #IV the third tract has separated from the triplex, rotating over itself and forming a single Hoogsteen bond with the top guanines (blue) of the first and second tracts, as described in Fig. 6*ε*. This basin represents the first relatively stable state of the system, corresponding to the time interval [0,3-0,7] *µ*s in Figs. 2 & 3. The fluctuations of these magnitudes in that time interval correspond to waving motions between the third and fourth guanine tracts.

Basins #V and #VI describe very similar situations. The last bond between the third tract and the remaining G-hairpin breaks. The third tract drifts away and pulls the second tract, breaking the G-hairpin and making it into the so called “cross-hairpin” (Fig. 6*η*). There is a slight difference between the two states, consisting in the relative position of tracts 3 and 4, which in basin #VI are a little closer due to a stacking interaction created by the top guanines (green) of both tracts (dashed circle in Fig. 10).

Basin #VII corresponds to the final state of the system: tracts 1 and 2 remain in a cross-hairpin, while the third and fourth remain close to each other, with two Hoogsteen bonds formed between them, almost constituting a complete hairpin. This configuration is very stable, with the presence of fluctuations due to the separation of the two blocks (see Figs. 2 and 3 from *t* = 0.8 *µ*s onwards).

With these results in mind, it becomes evident that tICA is able to provide us, upon selection of a proper value for the time window *τ*, a clear denaturation path. The states identified through the basins exhibit a high degree of dissimilarity, capturing the significant structural transformations and avoiding the oversampling of both transient and short-lived conformations.

In the figure, the links between states are bidirectional (two arrows) and weighted. They represent multiple transitions between the different basins with no information about a particular time sequence. This is due to the fact that, in principle, the system investigated lies in a dynamical equilibrium condition. This means that if the trajectories were long enough, they could be analyzed independently of time, with the occurrence of the same intermediates.

To study the stability of the basins found by tICA, the average escape time of each state can be calculated. These times reflect the average time the G-quadruplex remains in a given conformation before undergoing a transition. By analyzing escape times, insights into the stability and transition kinetics of different conformational states can be gained. The limitation of our analysis relies on the fact that we have a single unfolding trajectory, thus preventing us from performing broad statistics on the occupation of the different states. Nevertheless, we calculate the escape times by using two approaches: one based on Complex Markov Network self-loops and another analyzing the time spent in a given conformation, providing a an estimation of the free energy difference between the different basins. The detailed methodology as well as the results from these calculations can be found in the Supplementary Material; they reveal that the structure depicted in basin #IV has a larger lifetime than the rest, thus classifying it as a stable intermediate of the unfolding.

Additionally, the above analysis has been performed under a mesoscopic reduction of data, as previously explained. Nevertheless, the same tICA procedure can be applied without that intermediate step. In that case an even broader spectrum of states is resolved, with the resulting network still describing a clear denaturation path. This analysis is also included in the Supplementary Material.

#### 3.3.2 PCA results

Fig. 11 shows the network of states generated by the PCA procedure. It is evident that the interpretation of the basins becomes severely more challenging here when compared to the one produced by tICA. PCA aims to build coordinates containing the maximum variance possible which, when applied to the identification of unique basins of attraction, leads to the labeling of some equivalent states as different, overestimating the differences between equivalent configurations.

The initial state corresponds to the same starting conformation as in tICA, designated as basin #1 in Fig. <u>11</u>. Subsequently, basin #2 marks the point at which the 3^*′*^-end disengages from the primary structure, forming the G-triplex. Following this event, a number of tightly connected states is revealed (from basin #3 to #8). Basin #3 shows a slip-stranded G-triplex, with the fourth guanine tract away from the structure. Up to this point, the states analyzed by PCA are equivalent to those of tICA, namely basins #I, #II and #III.

Basin #4 shows a G-hairpin formed by the first and second tracts, with one of the guanines from the third tract in contact with the hairpin, while in basin #5 that contact has been broken, leaving a G-hairpin and two drifting guanine tracts (third and fourth).

The remaining basins (numbers #6, #7 and #8) in this heavily interconnected section of the network are classified as erroneously different by PCA respect to tICA. In particular, basins #6 and #7 depict a G-hairpin accompanied by an additional third guanine tract forming a single Hoogsteen bond. The difference between these conformations fundamentally lies in the relative orientation of bonds on the guanine columns forming the hairpin. Basin #8, on the other hand, is analogous to basin #5, represents a G-hairpin no longer in contact with the third tract. All these states (basins #5, #6, #7 and #8) identified by PCA are thus equivalent to basin #IV in tICA. Remarkably, the state identified by PCA only (basin #7) is then completely superfluous in describing the unfolding process because it is contained in basin #IV in tICA.

States #9, #10, and #11 and #12 show the final conformation of the system. They correspond to basins #V, #VI and #VII in tICA. They reveal the formation of transient and relatively short-lived bonds reminiscent of G-hairpins, resulting each of them in the division of the G-quadruplex into two well-separated guanine tracts: a cross-hairpin (tracts 1 and 2) and a disordered clump (tracts 3 and 4). However, while the three states found by tICA were distinguishable by the presence or absence of certain interactions (stacking between guanines in basin #VI and guanine-guanine bonds in basin #VII), in the case of PCA they are functionally equivalent: basins #10, #11 and #12 show the cross-hairpin in the first two tracts, along with an additional Hoogsteen bond between the third and fourth, while basin #9 presents an additional bond between them, with no stacking interaction being captured.

To conclude, our simulations reveal that PCA falls short in providing a minimal unfolding path akin to that offered by tICA, instead providing a multitude of interconnected states, some of them functionally equivalent.

Analogously as done with tICA analysis, the relative stability of the basins identified can be studied through the escape times, whose results can be found in the Supplementary Material. Furthermore, the PCA analysis without the mesoscopic reduction is also found in the Supplementary Material, leading to a higher number of basins with increased degeneration between them. They correspond to Supplementary Figures 6 & 7.

## 4 Discussion and conclusions

The study of unfolding pathways in biological systems is essential to understanding their stability and functional roles within their environments. In this work, we combined molecular dynamics simulations and multiple analytical techniques to explore the unfolding process of the human G-quadruplex in its parallel configuration (PDB: 1KF1). By characterizing key structural changes during unfolding, we have gained insight into the underlying factors that contribute to the stability and flexibility of this biologically relevant structure.

Replica Exchange Molecular Dynamics (REMD) simulations were employed to enhance conformational sampling. A distinct unfolding transition was observed in one of the eight replicas at temperatures near the denaturation point. Ion dynamics were found to play a fundamental role in this process: total ion loss appears to be a necessary step for unfolding, consistent with previous mechanical unfolding studies^17^. Systems that experienced only partial ion loss largely maintained their structural integrity, even reabsorbing ions from the environment and returning to their initial configurations, as evidenced in Fig. 4.

A mesoscopic reduction of the system coordinates^18^ allowed us to simplify the analysis while preserving key dynamics. In fact, comparison with the full-atom representation showed no significant differences in the observed trends (Supplementary Figs. 6–7). This opens the door for the creation and development of realistic coarse-grained models capable of reproducing the unfolding of these structures, similar to those applied to DNA chains^70^.

Dimensionality reduction techniques, specifically PCA and tICA, were then applied to the coarse-grained trajectories to identify the most relevant collective motions. The resulting output trajectories were used to construct Complex Markov Networks, revealing the main states and transitions that define the G-quadruplex unfolding pathway.

Our analysis of the simulations revealed a clear unfolding sequence. Firstly, partial strand separation is accompanied by ion loss from the central channel, allowing the unfolding to commence. Afterwards, one strand detaches, producing a G-triplex^22,61,62,71,72^, then rapidly reorganizing into a partially folded G-hairpin^22,67–69,71–74^, which from then onward remains as a fundamental structural motif, while exhibiting sometimes slip-stranding^65,66^ or accompanied by a cross-hairpin^69^. The transition sequence *parallel* G4 → triplex-like → hairpin, represents the dominant unfolding route captured in our simulations. Interestingly, the hairpin intermediate exhibits significantly longer persistence (∼0.4 *µ*s) when compared to the triplex intermediate (∼40 ns), indicating its relatively higher thermodynamic stability.

Experimental studies have also provided evidence for the existence of these kind of intermediates. Circular dichroism (CD) and single-molecule FRET (smFRET) measurements^63,64,74^ demonstrated that *antiparallel* and *hybrid* G4 structures, in both K^+^ and Na^+^ environments, can unfold through transient G-triplex and G-hairpin-like conformations. Although triplex-like intermediates are frequently observed, several single-molecule and spectroscopic studies have also reported alternative pathways that may bypass a well-defined triplex, involving partially folded or hairpin-like conformations instead^64,73–75^. Similar findings were obtained using UV-resonant Raman spectroscopy^76^, while time-resolved optical studies^77^ identified G-hairpin formation as a key early event in folding, consistent with the final conformations observed in our unfolding trajectories. Other techniques, including DNA origami^22,72^ and microfluidic mixing^71^, have revealed similar transient structures.

The main distinction between these experimental observations and our present results lies in the G4 topology. We focus here on the unfolding of a *parallel* G-quadruplex, in which all detected intermediates maintain parallel orientation. In contrast, the experimentally identified G-triplexes correspond to either *antiparallel* or *hybrid* structures^78^–direct evidence of parallel G-triplex intermediates remains elusive. Aznauryan *et al*.^75^ noted that fully parallel G4s are rarely stabilized under typical experimental conditions, complicating the observation of their unfolding pathways. Photochemical trapping studies^64,79^ also failed to identify long-lived parallel G-triplex species. The most plausible explanation is that parallel G-triplex intermediates possess extremely short lifetimes, making their detection experimentally challenging.

Our simulations are consistent with this interpretation. The distinct lifetimes registered for the intermediates can be coded into occupation probabilities and relative free energy differences. The calculations, though affected by the lack of a broad statistics, are detailed in the “Basin escape times” section of the Supplementary Material. The final state of the system (basin #VII in tICA, basins #9–#12 in PCA) includes a combination of G-hairpin and cross-hairpin features, confirming these as the most stable intermediates emerging from the parallel G4 structure.

Recent computational studies have further clarified the atomistic mechanisms of G-quadruplex folding and unfolding. Enhanced-sampling simulations reveal a highly rugged free-energy landscape populated by metastable triplex-like, hairpin, and slipped intermediates that interconvert through multiple pathways^50,80,81^. Kim *et al*.^81^ identified coexisting triplex-like and hairpin states during human telomeric G4 folding, while Pokorná *et al*.^50^ showed that parallel G4s can also fold via alternative routes that bypass a well-defined triplex. Janeček *et al*.^80^ likewise demonstrated multiple competing pathways in the folding of a parallel G4 from a single strand. Taken together, these results depict G4 unfolding/folding transitions as complex multi-pathway processes where specific linear series of events can probabilistically occur. Within this computational context, we believe our work contributes by uncovering one of the plausible unfolding routes of the parallel G-quadruplex, characterized by a sequential transition from the folded structure through triplex-like and hairpin intermediates.

Regarding the scope of our studies, the simulations were conducted at biologically relevant temperatures slightly above the melting point, thus ensuring that the observed conformations reflect thermally accessible states separated by realistic energy barriers. However, under these moderate thermal conditions, only one replica showed complete unfolding. The pathway observed reproduces intermediate steps that have been reported in both computational and experimental studies, the latter with different loops’ topologies, lending confidence to its biological relevance. Thus, more unfolding events are needed to uncover additional alternative unfolding pathways^50,80,81^, which would require extremely long trajectories in the conditions used in this work.

The sequence here investigated (5^*′*^-AGGGTTAGGGTTAGGGTTAGGG-3^*′*^) represents the minimal telomeric repeat capable of forming a parallel G-quadruplex. Extensions to longer sequences, as well as variations in loop size and rigidity^18,83,84^, cooperative interactions between adjacent G4 units^85,86^, molecular crowding^6,87,88^ and sequence constraints^89^, could influence both stability and kinetics. Besides all the many parameters that can be analyzed in the different physiological and biological contexts, our model focuses into the essential unfolding mechanisms followed by the parallel G-quadruplex. Similarly, the incorporation of other G4 topologies, such as antiparallel or hybrid, could lead to a more complete view of the G-quadruplex landscape. We have already made several attempts for both of these conformations, under the same conditions as the parallel simulations, without finding any significant results up to now.

With respect to the analysis methods, both tICA and PCA yielded Markov networks that effectively captured the relevant unfolding transitions. tICA proved particularly efficient in distinguishing kinetically distinct conformations, whereas PCA identified a larger number of states, several of which were structurally equivalent. Future studies may benefit from exploring non-linear dimensionality reduction and machine-learning-based techniques^90,91^, which could refine the description of conformational space, albeit at increased computational cost due to the need for larger ensembles of unfolding trajectories.

Finally, the last configurations obtained (basin #VII in tICA) suggest that refolding may not necessarily restore the original parallel topology. Instead, the unfolded state could evolve into alternative conformations such as antiparallel or hybrid G4s, as reported in other studies, showing a conformational switch between different G4 topologies, that warrants further exploration.

## Supporting information

Ion exits which trigger the unfolding

G4 Unfolding trajectory

## Author contributions statement

AF & FF conceived the project. AS-A conducted the simulations and the data analysis. AF, FF & AS-A wrote and reviewed the manuscript.

## Additional information

No competing interest is declared.

## Data Availability

All data generated or analyzed during this study are included in the published article. The structure used in the simulation corresponds to the human telomeric parallel G-quadruplex, found in the Protein Data Bank under the code 1KF1. The files of the structures identified in Figure 10 extracted from the simulations, in.pdb format, as well as all the gromacs parameter files.mdp, are available at the GitHub repository at the address: https://github.com/AlejandroSainzAgost/GromacsGQuadruplex.

