## Supplementary Material Document for "Telomeric G-quadruplex Intermediates unveiled by Complex Markov Network Analysis"

### 1 Uploaded Videos and .txt files.

- Video 1: Segment of the unfolding trajectory showing the exits of the ions which trigger the whole G4 unfolding.
- Video 2: G4 unfolding trajectory following the exit of the ions. It corresponds to the data plotted in Figure 2 in the interval from $t = 200$ ns to $t = 1200$ ns.
- The 7 files "Fig10_I-VII.txt" contain the structures plotted in Fig. 10, panels I to VII respectively, in .pdb format.

### 2 Mathematical description of PCA/tICA.

*PCA.* Consider a dataset **r**(t), encompassing information regarding the system's positions at various time instances, with the mean already subtracted from each coordinate. Our goal is to derive a linear combination of this input data that maximizes the variance.

$$\mathrm{Var}(\mathbf{r}v) = \mathrm{Var}\left(\sum_i v_i r_i\right) \text{ is maximal}$$

Where $v$ is the vector containing the coefficients of the linear combination that we aim to obtain. From the properties of the variance, this expression can be rewritten as

$$\begin{aligned}&\mathrm{Var}(Ra) = v_1^2\mathrm{Var}(x_1) + v_2^2\mathrm{Var}(x_2) + \\ &2v_1v_2\mathrm{Covar}(x_1x_2) + \cdots = v^T\mathbf{C}(0)v\end{aligned}$$

Where $C(0)$ corresponds to the covariance matrix of **r**(t), the 0 denoting that the covariance is calculated between the values of **r(t)** at the same time instances. This notation is relevant, and will come in handy later. The problem of obtaining the optimal values for $v$ can be solved employing Lagrange multipliers. Introducing the restriction that $v$ must be a unit vector, i.e. $v^T v = 1$, we have:

$$v^T\mathbf{C}(0)v - \lambda(v^T v - 1) = 0 \overset{\partial/\partial v}{\Longrightarrow} \mathbf{C}(0)v = \lambda v \tag{1}$$

Thus, the problem of obtaining the values of $v$ is, in fact, an eigenvalue problem of the covariance matrix, in which $v$ is the eigenvector and $\lambda$ the eigenvalue. The latter also coincides with the variance of the data projected into the direction marked by $v$. Since the covariance matrix is symmetric and real, all of its eigenvalues will be real, and we can extend this procedure to obtain $N$ vectors. Generalising:

$$\mathbf{C}(0)\mathbf{V} = \mathbf{\Lambda}\mathbf{V} \tag{2}$$

Where $V$ is the matrix containing the eigenvectors $v_i$ by columns, and $\Lambda$ is the diagonal eigenvalue matrix. Then, projecting our input data into the new coordinates, we obtain the *Principal Components*: $\mathbf{x}^T(t) = \mathbf{r}^T(t)V$.

*tICA.* Given a value of $\tau$, we want the autocorrelation of the projection between $t$ and $t+\tau$ to be maximal, defined as

$$\begin{aligned}\mathrm{Autocorr}(\tau, \mathbf{r}w) = \langle \mathbf{r}w(t)\mathbf{r}w(t+\tau)\rangle &= w_1^2\langle r_1(t)r_1(t+\tau)\rangle \\ &+ w_2^2\langle r_2(t)r_2(t+\tau)\rangle + w_1w_2\langle r_1(t)r_2(t+\tau)\rangle \\ &+ w_1w_2\langle r_2(t)r_1(t+\tau)\rangle + \cdots = w^T C(\tau)w\end{aligned}$$

Where $C(\tau)$ is the time-lagged correlation matrix, whose element $ij$ is given by

$$C_{ij}(\tau) = \frac{1}{N-\tau-1}\sum_{t}^{N-\tau} r_i(t)r_j(t+\tau)$$

In order to eliminate the effect of the variance in the eigenvalues of the system and be able to select modes based on timescale alone, our restriction is to force our coordinates to have unit variance: $w^T C(0)w = 1$, leaving

$$w^T \mathbf{C}(\tau)w - \gamma(w^T \mathbf{C}(0)w - 1) = 0 \tag{3}$$

Corresponding to the generalized eigenvalue problem of

$$\mathbf{C}(\tau)\mathbf{W} = \mathbf{C}(0)\mathbf{W}\boldsymbol{\Gamma} \tag{4}$$

Where $\Gamma$ is the diagonal eigenvalue matrix, and $W$ the eigenvectors matrix, with each of the different $w$ vectors as columns.

The general Eq. (4) is typically not solvable due to the small value of the determinant of the matrices involved, leading to numerical errors in the calculations. The AMUSE algorithm[2] is typically used in its place, consisting of several steps:

1. Subtract the mean from the data: $\mathbf{r}'(t) = \mathbf{r}(t) - \langle \mathbf{r}(t) \rangle_t$.
2. Compute the PCA of the data, and project it into the obtained eigenvectors:

$$\mathbf{C}(0)\mathbf{V} = \boldsymbol{\Lambda}\mathbf{V} \;\rightarrow\; \mathbf{x}^T = \mathbf{r}^T V \tag{5}$$

3. Normalise the newly obtained principal components by their corresponding eigenvalue: $\mathbf{x}'(t) = \Lambda^{-1}\mathbf{x}(t)$
4. Build the time-lagged covariance matrix of **x**'(t), symmetrize and diagonalise it:

$$\mathbf{C}(\tau)\mathbf{U} = \boldsymbol{\Gamma}\mathbf{U} \tag{6}$$

   Note that the eigenvalues obtained in this step coincide with those in (4), but the eigenvectors do not. Since the latter are the coordinates of $W$ in the space defined by $V\Lambda^{-1}$, it can be verified that $\mathbf{W} = \mathbf{V}\boldsymbol{\Lambda}^{-1}\mathbf{U}$.

5. Project $\mathbf{x}'$ onto these new eigenvectors, $\mathbf{y}^T = \mathbf{x}'^T\mathbf{U}$. This is the projection of our original data into the tICA-defined coordinate system.

Identically to PCA, a few components of $\mathbf{y}(t)$, the ones corresponding to the larger timescales, can be employed to study the whole system.

### 3 Simulation figures by replica ID.

Supplementary Figs. 1 & 2 contain the evolution of the RMSD and $R_g$ respectively, for the different replicas. These results, as the ones in the main text, are shown both using the full G-quadruplex in the calculation (blue lines) and using only the G-tetrads, excluding the loops (orange lines). The grey vertical dashed lines indicate successful exchanges between them, with the destination replica shown above them.

The unfolding event characterized by a sudden increase in RMSD and $R_g$ begins in Rep. 4, corresponding to Sys. 5 after the exchange at 0.2 $\mu$s. The high values in the graph remain up to 1.2 $\mu$s, in which an exchange between Reps. 4 & 3 takes place, leading to the abrupt increases in the latter. This information could be mistakenly interpreted as an additional unfolding event in Rep. 3, thus the choice in presentation in Figs. 2 & 3 of the main text.

### 4 Basin metastability.

Each of the basins found during our analysis corresponds to a significant conformation the system adopts at some point in its unfolding trajectory. The stability of these conformations is a matter of interest, allowing us to classify them into either stable intermediates or fast-lived (although necessary) transition states, contributing information to the energy landscape of G-quadruplexes.

One possible measure to study the relative stability of these basins is the average escape time, *i.e.* the typical time the G-quadruplex remains in a particular conformation before transitioning into another state. The escape time $t_{\text{esc}}$ can be estimated

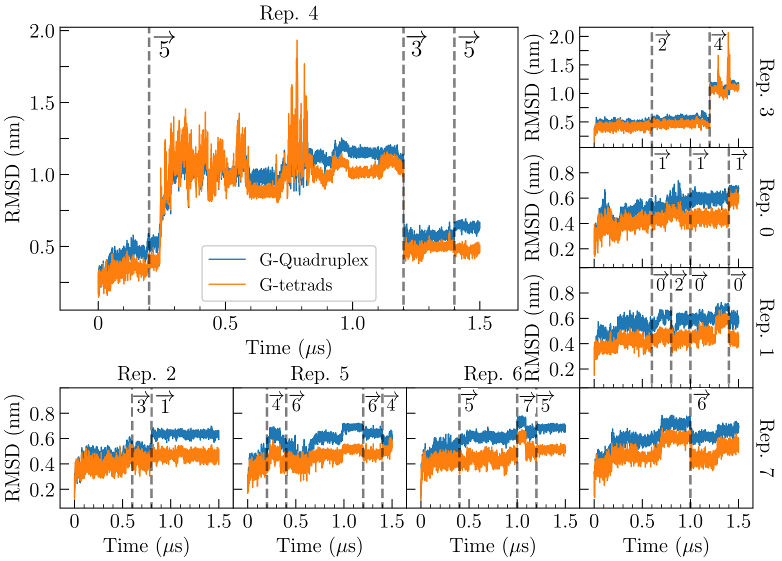


**Supplementary figure 1.** RMSD calculated over the different replicas (Rep. ID in each graph) of the parallel G-quadruplexes. In blue, the RMSD of the whole structure. In orange, the RMSD taking into account only the guanines forming the planar arrangement. The vertical dashed lines correspond to successful exchanges, with their destination replica shown on their right under an arrow. Rep. 4, since its the one used for the analysis, has been enhanced.

using the self-loop probabilities associated with each basin, whose weight represents the probability of remaining in the same state at the next timestep,

$$t_{\text{esc}} = \frac{\Delta t}{1 - P_{\alpha\alpha}} \tag{7}$$

where $\Delta t = 20\,\text{ps}$ is the time resolution of the trajectory used to construct the Complex Markov Network, and $P_{\alpha\alpha}$ represents the self-loop probability of basin $\alpha$.

Another possibility for studying stability of states relies on computing the times the trajectory remains into a given basin along the simulation, named $t_{\text{TRAJ}}$.

Finally, from the residence time in the different basins $t_{\text{TRAJ}}$ we calculated the relative weight of these states along their trajectory, $P_\alpha$. The free energy difference of each basin to a reference state (chosen as the most populated basin) can be computed as

$$\frac{\Delta F_\alpha}{k_B T} = -\log\left(\frac{P_\alpha}{P_{\text{ref}}}\right) \tag{8}$$

*tICA.* The results of all the above described stability measurements for tICA ($\tau = 0.7$ ns) can be found in the table on the top of supplementary figure 3. All quantities show relatively high values on basins #I, #IV and #VII. This is in line with our results: the system remains during an initial time interval in the original conformation (basin #I) before undergoing unfolding, then it lasts a long time in the final conformation (basin #VII). Basin #IV reveals to be a stable intermediate, surviving for a large portion of the simulation.

These observations are further confirmed by the figure on the bottom panel of supplementary figure 3, which shows the evolution of the system over time in terms of basins, showing the time spent in each of them.

The difference between the values of $t_{\text{esc}}$, which underestimate the stability of the basins, and the rest of the measurements are fundamentally linked to a lack of statistical data. Changes between basins are not numerous, and their values tend to be concentrated around low times, heavily affecting the self-loops.

Additionally, basins #I and #VII are affected by our choice of the beginning and end points of the trajectory. Since our analysis starts after the replica exchange at 0.2 $\mu$s and unfolding begins around 0.25 $\mu$s, the stability of the initial state of the trajectory is underestimated. The same happens for the final state of the trajectory, although to a lesser extent: trajectory analysis ends at 1.2 $\mu$s, but the system remains in that state an additional 0.3 $\mu$s, not taken into account due to the replica exchange taking place in between.

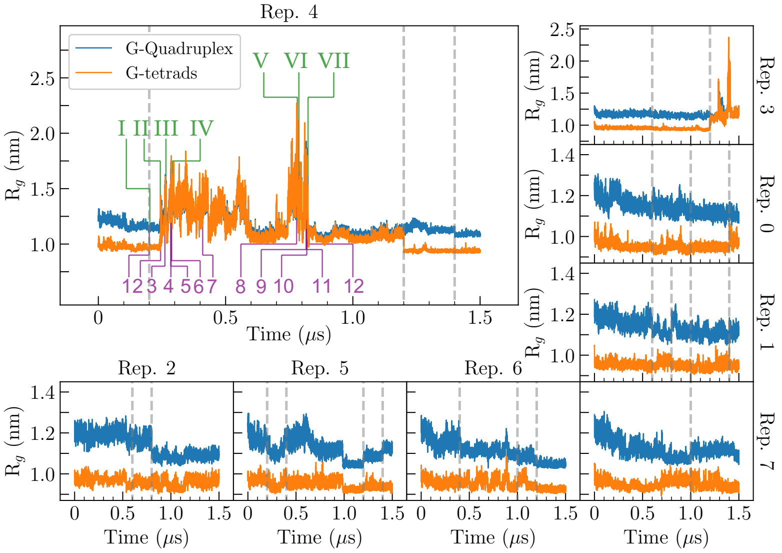


**Supplementary figure 2.** Radius of gyration computed over the different replicas of the parallel G-quadruplex. In blue, the $R_g$ of the whole structure. In orange, the $R_g$ for the G-tetrads. The fourth replica has been enlarged, since it is the one considered for posterior analysis. The vertical grey dashed lines correspond to successful exchanges, with their destination replica shown with a number under an arrow on the right of the lines. The purple and green lines correspond to the states identified by PCA and tICA respectively, and are further explained in later paragraphs.

*PCA.* The results of escape time $t_{esc}$, time spent during the trajectory $t_{TRAJ}$, relative weight $P_\alpha$ and relative free energy difference $\Delta F/k_B T$ for PCA can be found in the table on the left panel of supplementary figure 4. Similarly to tICA, all stability measurements exhibit relatively high values on some basins. Particularly, basin #1, corresponding to the initial state of the simulations, basins #7 or #8, corresponding to the stable intermediate (basin #IV for tICA) and basins #10 or #12, the final state of the system (basin #VII for tICA).

However, these results are misleading. PCA, as explained in the Section 3.3 of the main text, tends to identify different states that are in reality functionally equivalent. Basins #9 to #12 (final state of the trajectory, basin #VII for tICA) correspond to the same conformation but are classified as different by PCA. This leads to 4 interconnected states, that have reduced self-loops and lifetimes, leading to a decrease in $t_{CMN}$ and $t_{traj}$ which, in terms of stability, leads to one of the states being classified as extremely stable (basin #12), while the remaining are underestimated. This same phenomenon happens with basins #5, #6, #7 and #8, corresponding to basin #IV in tICA.

The disparity in stability and time spent in each state between equivalent basins can also be seen in the figure on the right panel of Fig. 4, where the sum of basins #5, #6, #7 and #8 would lead to basin #IV in tICA, with basins #6 and #8 being practically not populated. As for the final state of the trajectory, the graph shows how basins #9, #10, #11 are irrelevant for the description, their difference with basin #12 being an artifact produced by PCA.

### 5 Relation between the states skipped and the resulting networks.

As commented previously in Section 3.3 of the main text, a valid method to reduce the final number of basins identified in the construction of the Complex Markov Network consist on, instead of taking every frame of the trajectory to build such network, only processing one of every $X$ frames.

Note that this step may take place only after the application of the dimensionality reducing algorithms in order to have a common ground on which to compare the results of the final network, and to prevent the creation of artificial coordinates not representative of the system at hand.

The idea of the method is that, when changing the number of frames read, we may avoid the oversampling of certain regions not of interest for the unfolding and/or conformational changes of the system (i.e. regions of relatively long-lived transient states) and thus potentially enhance the presence of states that, while short-lived, may be of importance to the observed phenomena. Obviously, a risk one can incur into when applying this technique is to skip more steps than necessary; this eventually leads to the detection of a single basin of attraction. Supplementary fig. 5 contains the number of detected basins as a function of the number of skipped states in the trajectory for both PCA and tICA ($\tau = 0.7$ ns).

Additionally, for both the PCA and tICA methods the final single basin detected corresponds to a native conformation. This

| Basin | #I | #II | #III | #IV | #V | #VI | #VII |
|---|---|---|---|---|---|---|---|
| $t_{esc}$(ns) | 1000.0 | 0.17 | 0.75 | 1.23 | 0.50 | 0.97 | 24.69 |
| $t_{TRAJ}$(ns) | 44.24 | 21.1 | 21.46 | 492.1 | 36.82 | 6.8 | 377.48 |
| $P_\alpha$ | 0.044 | 0.021 | 0.022 | 0.492 | 0.037 | 0.007 | 0.377 |
| $\Delta F/k_B T$ | 2.409 | 3.149 | 3.132 | 0.0 | 2.593 | 4.282 | 0.265 |

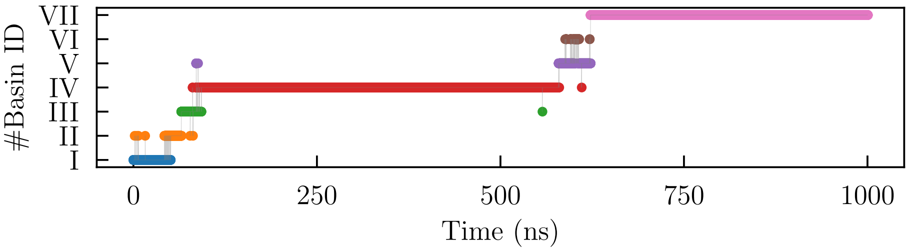


**Supplementary figure 3.** Table: different magnitudes for the basins detected by tICA with $\tau = 0.7$ ns. In the first row, escape times $t_{CMN}$ calculated with Eq. 7. Second row, total time spent in each basin during the trajectory $t_{\mathrm{TRAJ}}$. Third row, relative weight of each basin $P_\alpha$. Fourth row, relative energy difference respect to the most stable basin (largest $P_\alpha$), calculated with Eq. 8. Figure: trajectory of the G-Quadruplex during its unfolding expressed in terms of basins.

| Basin | #1 | #2 | #3 | #4 | #5 | #6 |
|---|---|---|---|---|---|---|
| $t_{CMN}$(ns) | 63.50 | 0.09 | 0.09 | 0.08 | 0.50 | 0.16 |
| $t_{TRAJ}$(ns) | 44.54 | 28.02 | 15.36 | 22.22 | 312.74 | 7.44 |
| $P_\alpha$ | 0.0148 | 0.0093 | 0.0051 | 0.0074 | 0.1042 | 0.0025 |
| $\Delta F/k_B T$ | 3.983 | 4.447 | 5.048 | 4.679 | 2.034 | 5.773 |

| Basin | #7 | #8 | #9 | #10 | #11 | #12 |
|---|---|---|---|---|---|---|
| $t_{CMN}$(ns) | 8.89 | 0.18 | 0.23 | 0.62 | 0.15 | 0.14 |
| $t_{TRAJ}$(ns) | 172.06 | 0.92 | 2.7 | 1.18 | 1.64 | 391.72 |
| $P_\alpha$ | 0.0574 | 0.0003 | 0.0005 | 0.0003 | 0.0004 | 0.7972 |
| $\Delta F/k_B T$ | 2.632 | 7.780 | 7.285 | 7.863 | 7.614 | 0.0 |

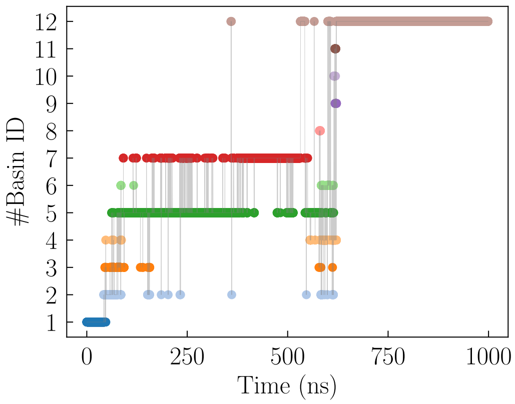


**Supplementary figure 4.** On the left, table with different magnitudes for the basins detected by PCA. In the first row, escape times $t_{esc}$ calculated with Eq. 7. Second row, total time spent in each basin during the trajectory $t_{\mathrm{TRAJ}}$. Third row, relative weight of each basin $P_\alpha$. Fourth row, relative energy difference respect to the most stable basin (Eq. 8), with the reference taken as the basin with highest $P_\alpha$. On the right, the trajectory of the G-Quadruplex during its unfolding expressed in terms of basin's occupation.

is due to the fact that, when we skip enough steps, all of the subsequent recorded nodes of the network are connected to the first step of the trajectory, which is a native conformation.

However, analyzing low values of the number of skipped frames for both methods quickly reveals that the effect of this parameter is the opposite: while we do indeed reduce the number of basins detected, the native state of the system is heavily over-represented. This may be due to how the stochastic steepest descent works, making connections between unrelated nodes and grouping them under a single representative basin, corresponding to the native state. After seeing this results, no states were skipped in the construction of any of the networks.

### 6 Effect of the coarse-graining of the trajectories.

In our analysis method, the trajectories of the system obtained from the all-atom simulations in Gromacs are converted into a reduced number of coordinates, following the description of the G-quadruplex mesoscopic model[3]. However, given how later either PCA or tICA are applied onto said trajectories, the mesoscopization of the coordinates is not a necessary stage, but rather one we introduced as to reduce the possible influence of the relative orientations of the side chains onto the final results of PCA and tICA. Here we present the final networks for both methods without the mesoscopic discretization applied. The networks contained in Supplementary figs. 6 & 7 represent the results of the analysis without the application of the mesoscopic

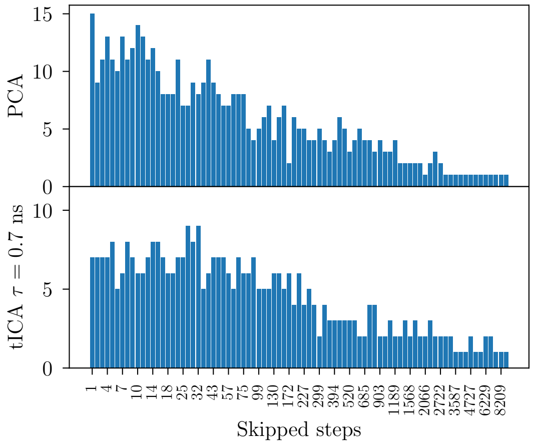


**Supplementary figure 5.** Number of basins in the final Complex Markov Network as a function of the frames skipped in its construction for both PCA (top) and tICA ($\tau = 0.7$ ns, bottom). As stated in the text, both methods eventually detect a single basin. The top limit for the number of skipped frames is set arbitrarily; a single frame was detected from then onwards.

coarse-graining.

#### 6.1 tICA ($\tau = 0.7$ ns)

The network uncovered by tICA is essentially the same as the one encountered in the main text: starting from the native folded conformation (basin #I) we observe the formation of the G-triplex, with a rogue guanine chain separated from the rest (basins #II and #III). That G-triplex is further split, leading to a G-hairpin core and drifting guanines. However, different to the results after coarse-graining, here we have the coexistence of 3 interconnected states depicting the formation of said G-hairpin.

Basin #IV describes the G-hairpin with a single Hoogsteen bond with the rotated guanine tract 3, corresponding to the top plane, basin #V describes the G-hairpin with that Hoogsteen bond already broken, and finally, basin #VI shows the rotation of one of the guanine chains forming the G-hairpin into a situation similar to the one encountered in the previous state. Basin #V is only connected with basin #VI since they are equivalent between each other.

After that, the network returns to more familiar territory; nodes #I and #II show the collapse of the G-quadruplex into a disordered lump after its complete denaturation, corresponding to the final state of the recorded trajectories.

Thus, in the case of tICA, no notable differences between the networks are recorded, save for the interconnection of the G-hairpin like states, which is not contemplated after coarse-graining. However, the links between nodes #V and #VII seem rather fictitious; one would expect to first break the remaining Hoogsteen bonds like the situation found in #IV, and may reflect the effect that considering the side chains for the analysis may have in our system. All nodes identified in this network were around the same time intervals than the ones detected before by tICA.

#### 6.2 PCA.

For PCA, the network increases significantly in complexity, having 17 different basins well interconnected between them. Supplementary fig. 7 contains the pertaining network for this case.

This network fundamentally contains the same unfolding process (although with more nodes present) found in previous paragraphs. However, given the degree of interconection between the basins, trying to extract an evolution over time of the state of system is extremely difficult, even more so than in the previous PCA network.

The evolution of the system begins with the native state at node 1. Afterwards, basins 2, 3, 4 and 5 describe the transformation of the native state into the G-triplex, with every one of those nodes progressively describing the breaking of said G-triplex. Basin 17 is the final stage of that structure, presenting a slip-stranded hairpin and a cross-hairpin.

Subsequently basins 7, 8, 9 and 10 show the dynamics of the G-hairpin. The first three show the system of two guanine columns in various states of arrangement respect of the angle between bonds and the state of the rogue chains, while the last one shows the moment in which the hairpin is broken and one chain has rotated over itself, forming, occasionally, individual Hoogsteen bonds.

The rest of the basins fall in some category in between those two, representing transient states found in the system. An example of this is node 11, which represents a hairpin with a single bond with another guanine, transitioning into a full hairpin.

This confirms that for PCA, which already had problems elucidating a clear unfolding path when the mesoscopic reduction was applied, only leads to worse results when omitting this step. No additional relevant intermediate states were revealed

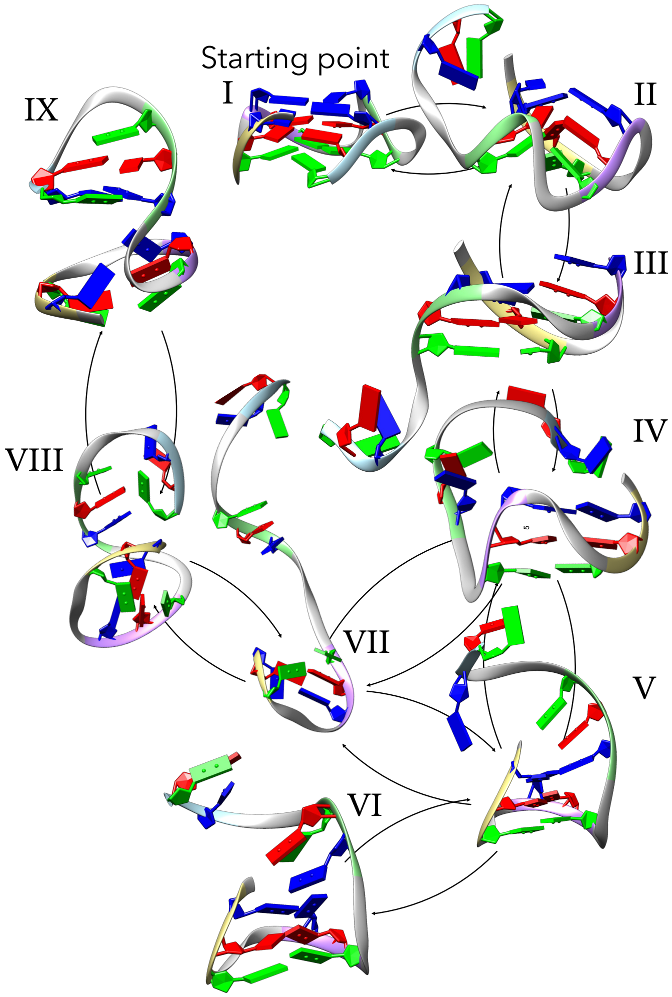


**Supplementary figure 6.** Network of basins found for the tICA method, with $\tau = 0.7$ ns, without the mesoscopic discretization of the trajectories. Note that the IDs on the nodes do not correspond to the ones shown in the main text.

respect to previous realizations of both PCA and tICA with the coarse-graining applied.

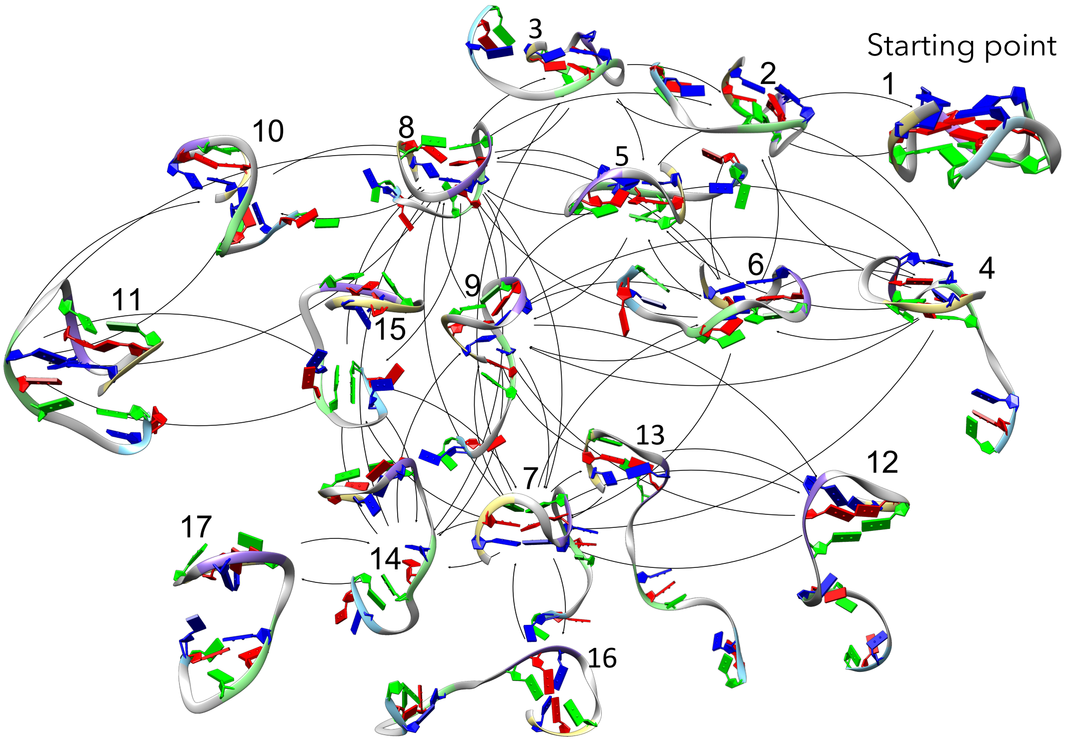


**Supplementary figure 7.** Network of basins found for the PCA method without the mesoscopic discretization of the trajectories. Note that the IDs on the nodes do not correspond to the ones shown in the main text.

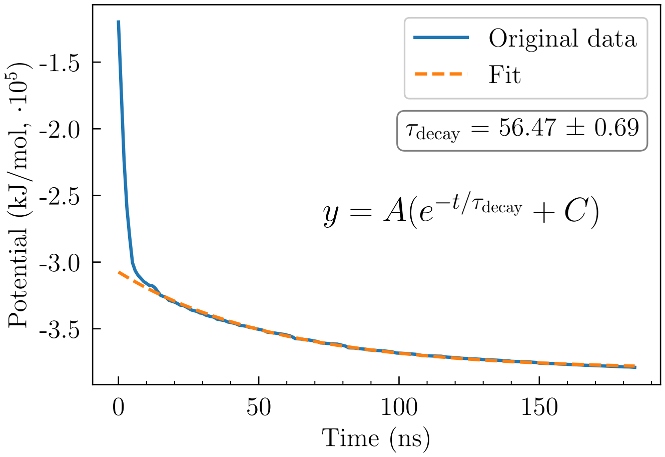


**Supplementary figure 8.** Evolution of the potential energy of the lowest temperature replica (343 K). In blue, the original data. In orange, the fit to an exponential function, whose expression and decay rate $\tau_{\text{decay}}$ are contained in the figure.
